# An Artificial Cofactor catalyzing the Baylis-Hillman Reaction using Designed Streptavidin as Protein Host

**DOI:** 10.1101/2020.03.05.978098

**Authors:** Horst Lechner, Vincent R. Emann, M. Breuning, Birte Höcker

## Abstract

An artificial cofactor based on an organocatalyst embedded in a protein was used to conduct the Baylis-Hillman reaction in a buffered system. As protein host we chose streptavidin, since it can be easily crystallized and thereby supports the design process. The protein host around the cofactor was rationally designed based on high-resolution crystal structures obtained after each variation of the amino acid sequence. Additionally, DFT-calculated intermediates and transition states were used to rationalize activity. Finally, repeated cycles of structure determination and redesign led to a system with 24 to 35-fold increased activity over the bare cofactor and to the most active proteinogenic catalyst for the Baylis-Hillman reaction known today.

---

The design of proteins that display new catalytic activities is still a major challenge. Although several successful examples were reported^[1–3]^, these *de-novo* cases are limited to a small set of reactions, such as the Kemp elimination^[4,5]^ and the Retro-aldol reaction^[6]^ as well as a bimolecular Diels-Alder reaction.^[7]^ All the initial designs provided a sufficient starting point but had to be strongly improved through directed evolution to show reasonable rate enhancement.^[8–10]^

Of all the natural enzyme catalyzed reactions half of them require cofactor(s) as part of their catalytic machinery.^[11]^ Hence it might not be surprising that far more new protein-based catalysts were reported using artificial (metal-based) cofactors embedded in host-proteins.^[12–16]^ These cofactors bear the advantage of intrinsic activity, which is usually low without surrounding protein. The protein can provide a different environment for the catalysis – increasing in most cases activity and maybe even facilitating (stereo)selective reactions.

There are a number of examples using streptavidin as host for these cofactors, since its natural ligand biotin binds strongly (K_d_ ~ 10^−15^ M) and is easy to modify with catalysts at the carboxylic acid group. The location of the catalyst is a shallow cavity at the surface of the protein. Streptavidin is a known thermo- and solvent stable protein, which is another advantage for developing new catalysts.

Contrary to metal-based cofactors, organocatalysts are rarely used in this context. There are only few examples: proline^[17]^ and an imidazolium salt^[18]^ were used in wildtype (*wt)*-Streptavidin as host for Michael additions and Aldol-reactions, respectively.

No enzyme is known to naturally catalyze the Baylis-Hillman reaction^[19,20]^, a very versatile C-C bond forming reaction for the production of various compounds^[21]^ and intermediates of APIs^[22]^. This reaction provides in an atom-economic way a very functionalized product. There were attempts to design a *de-novo* enzyme but with only minor success.^[23]^ Nevertheless it was recognized that this reaction can be catalyzed using nucleophilic amine catalysts in aqueous systems, although with low rates and high catalyst concentrations. (Imid)azoles^[24,25]^, DABCO^[26,27]^, 3-hydroxyquinuclidine^[27]^ and 4-dimethylaminopyridine (DMAP)^[27]^ were reported as catalysts for this purpose.

We chose a derivative of the latter to be used as an artificial cofactor due to its low steric expansion, and supposedly easier to include in a protein scaffold. Thus, the DMAP derivative 4-(4-aminopiperidino)pyridine was used to synthesize the artificial cofactor **4** (Figure 1, B).

**Figure 1.**
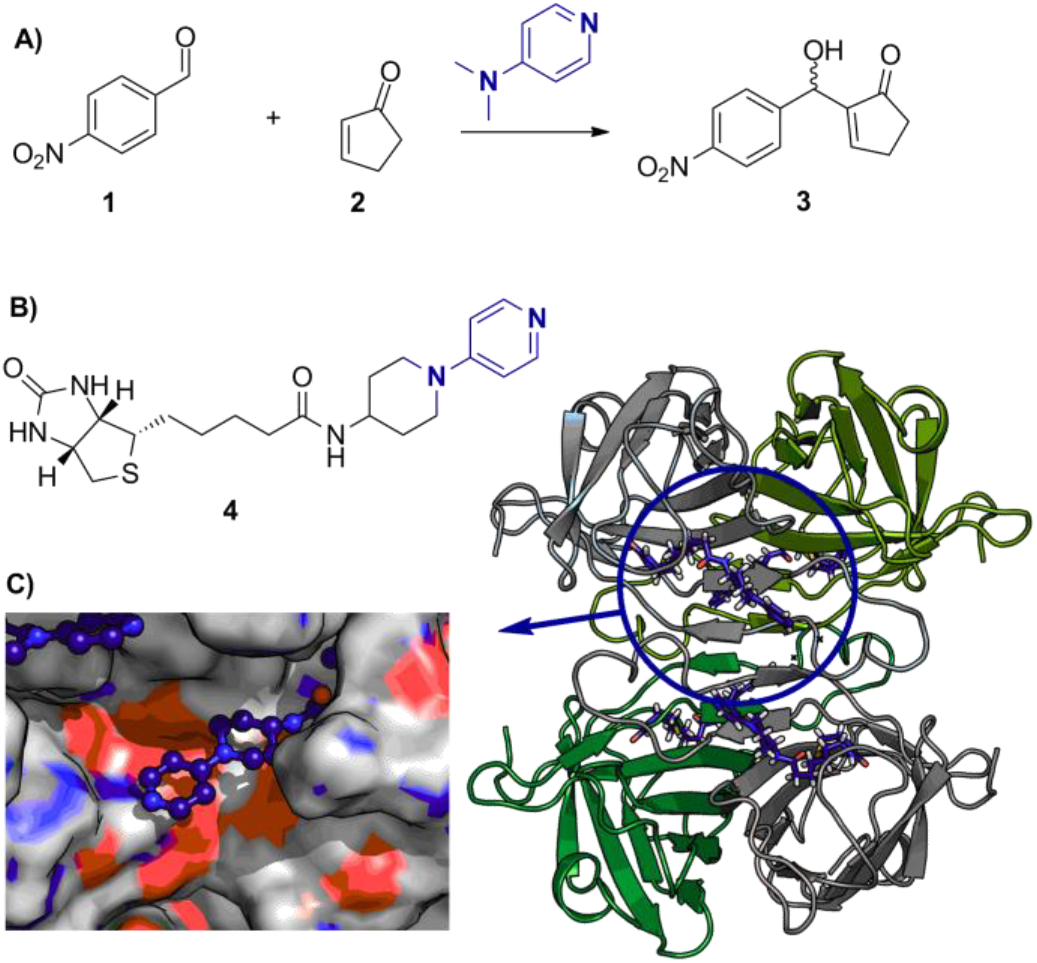
Concept of the artificial cofactor - protein host system as catalyzing the Baylis-Hillman reaction. **A)** DMAP used as catalyst for the reaction of an activated alkene **1** with an aldehyde **2** to yield the racemic alcohol **3, B)** Artificial cofactor **4** consisting of biotinylated DMAP-derivative, **C)** The crystal structure of the tetrameric *wt*-streptavidin harboring **4** and a closeup of the binding site of **4**.

As protein host streptavidin was used due to its aforementioned advantages. The wt-protein was expressed, purified and after adding **4**, the activity for the reaction tested. *p*-Nitrobenzaldehyde **1** and cyclopentenone **2** were used as model substrates (Figure 1, A). No activity above the activity seen in a blank reaction without cofactor could be observed. Consequently, the x-ray structure was solved (Figure 1C, PDB: 6T1E) to reveal the causes. It clearly visualized that **4** is placed nicely at the entrance of the biotin-binding pocket. The density of the ligand is very well resolved (see Composite Omit maps, Figure S1). As depicted in Figure 1, C and Figure 2, A, the catalyst is sterically hindered by the sidechains of Q114 and R121. Especially position 121 is known to influence the outcome of metal-based catalysts to a great extent.^[28]^ Thus, the sterically demanding Q114 was changed to A or T, and R121 and S112 were substituted by A as well. Additionally, the variant S112A, R121A was tested, but none of these variants showed activity higher than the background (Table 1, Entry 4 and 5). Exemplarily the activity of streptavidin S112A Q114A R121A is shown in Table 1, Entry 6. The derived crystal structures of these variants revealed that **4** was moved to a position close to the amide carbonyl oxygen of A121, leading to a non-accessible nucleophilic nitrogen atom (Figure 2, B yellow). An attempt of introducing Y as a bulkier amino acid at position L124 behind the catalyst to push it forward was not successful either (Figure 2, B pink). The next residue that caught our attention was S112 beneath the catalyst. By increasing its size from A to its original S and further increasing this residue to M, F and I we reasoned it might force the catalyst in a better position. The variant harboring S112 led to no activity and a similar structure was yielded (PDB: 6T30) as in round 1 and 2 of our design approaches. In contrast, all variants harboring sterically more demanding residues at position 112 (I, F, M) displayed activity (Table 1, Entries 7-11). Our reasoning was supported by the x-ray structures of these variants (Figure 2, C) where the position of the nucleophilic nitrogen of **4** superimposes with the one from the initial structure. The S112I variant displayed the highest activity among the three variants (Table 1, Entry 9). The S112F variant, although structurally very similar to S112I (Figure 2, C) catalyzed roughly half of the conversion as the S112I variant (Table 1, Entry 10). Reasons for this might be the π-π interaction of F112 and the catalyst, keeping it too tightly together as explained below. S112M (PDB: 6T2Z) led to two different conformations of **4** (Figure S1) and very low activities (Table 1, Entry 11). While all these constructs are active, they however lack enantioselectivity.

**Table 1.** Activities and selectivities of catalysts and selected streptavidin constructs for the model substrates **1** + **2**.

| Entry | Catalyst | Mol%<br>Cat. | Reaction<br>time [h] | Conv.<br>[%]<sup>[a]</sup> | e.e.<br>[%]<sup>[a]</sup> |
| --- | --- | --- | --- | --- | --- |
| 1 | - | - | 48 | 0 |  |
| 2 | DMAP | 2 | 48 | 3 |  |
| 3 | **4** | 2 | 48 | 1 |  |
| 4 | Streptavidin<sup>[b]</sup> |  | 24 | 6 |  |
| 5 | Streptavidin |  | 48 | 11 | <5 |
| 6 | **S112A** Q114A R121A<sup>[b]</sup> + **4** | 1 | 24 | 8 |  |
| 7 | **S112I** Q114A R121A<sup>[b]</sup> L124Y + **4** | 1 | 24 | 13 |  |
| 8 | **S112I** Q114A R121A L124Y + **4** | 2 | 24 | 16 |  |
| 9 | **S112I** Q114A R121A L124Y + **4** | 2 | 48 | 35 | <5 |
| 10 | **S112F** Q114A R121A L124Y + **4** | 2 | 48 | 17 | <5 |
| 11 | **S112M** Q114A R121A L124Y + **4** | 2 | 48 | 12 | <5 |
| 12 | K49N **S112I** Q114A R121A L124Y **H127Q** |  | 48 | 9 |  |
| 13 | K49N **S112I** Q114A R121A L124Y **H127Q** + **4** | 2 | 48 | 23 |  |

Reaction conditions: HEPES buffer (10 mM, pH 7.0), streptavidin (2.6 mol% of monomer), DMSO (20 vol%), 4-nitrobenzaldehyde (50 mM), cyclopentenone (100 mM), 30 °C, orbital shaking 1000 rpm. [a] Determined by HPLC in duplicates [b] 1.3 mol% of monomer.

**Figure 2.**
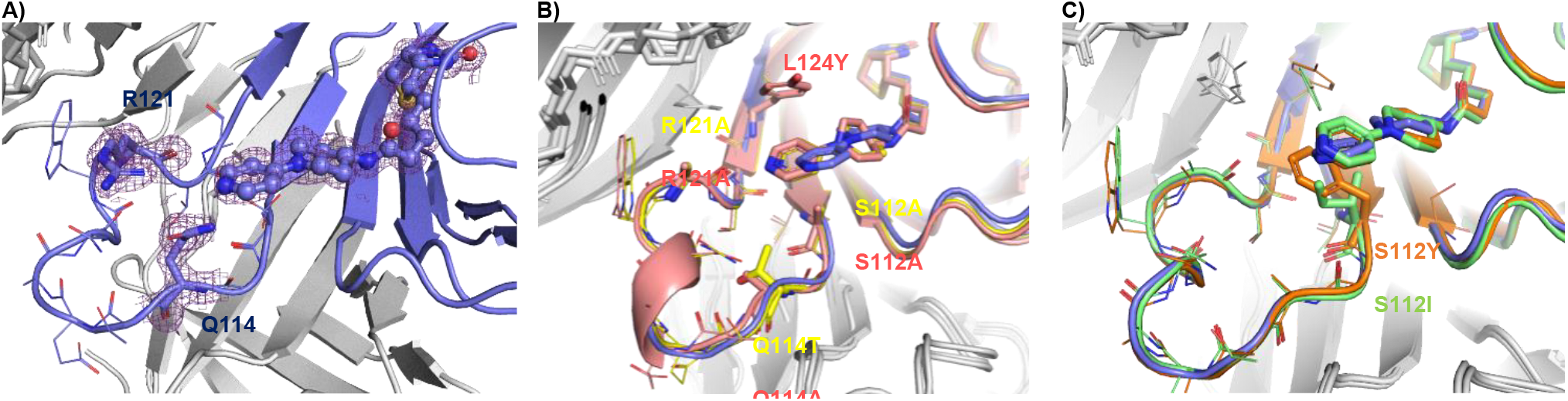
Crystal structures obtained during the course of catalysts development. Chain A of corresponding structure is colored, chain B, C and D are in grey. **A)** *wt*-streptavidin (violet; PDB: 6T1E) displaying **4**, blocked by residues Q114 and R121. Density obtained from the 2mFo-DFc map. **B)** First and second round of design: S112A, Q114T and R121A (yellow, PDB: 6T1G) as well as S112A, Q114A, R121A, L124Y (pink, PDB: 6T2Y). Both variants have space at the original positions of the catalyst, but the catalyst moved tilted backwards in a non-accessible position. **C)** Third round of design where additional to S112A, Q114A, R121A and L124Y position S112 was changed to F (orange, PDB: 6T31) and I (green, PDB: 6T32) leading to a good position of the catalyst within the protein environment.

Two major questions arise with these results in hand: What is the cause for the high “background” reaction displayed by streptavidin alone? And why does the active protein-cofactor system lack enantioselectivity?

We want to address the question regarding the high background first. In this regard it is worth mentioning that others^[29]^ reported some proteins at high concentrations (30 mg/ml), such as the carrier protein BSA, being able to catalyze the Baylis-Hillman reaction. The mechanism of catalysis was however not elucidated further. We considered histidine residues at the surface of streptavidin as the cause for catalysis, since it was described that (imid)azoles^[24,30]^ can serve as catalysts under certain conditions. Streptavidin has two histidine residues per monomer located at the surface (H87, H127) (Figure S2). H127 is known to be tolerant against mutations.^[31]^ Therefore we created a double mutant of the S112I variant with a H127Q and K49N exchange. The latter exchange removes a lysine close to the catalyst, which could also have an effect on the outcome of the reaction. While the conversions dropped somewhat (Table 1, Entry 12 and 13), they were still comparable to the results obtained without these mutations. Thus, H127 has minor importance for the background reaction. Exchange of H87 was already recognized to disturb the integrity of the protein.^[32]^ After careful inspection of the structures we suspect that this histidine forms a hydrogen bond to an aspartic acid of a neighboring chain (D61) keeping loops of neighboring subunits of streptavidin together and creating a “catalytic diad” by activating the histidine, which would explain the outcome of our experiments. Our efforts to change this H87 to a D, N or S unfortunately lead to misfolded protein, only the double mutant H87Y and D61I could be refolded with very low yields and diminished biotin binding ability. As a consequence, this variant could not be characterized further.

Next we addressed the issue of enantioselectivity in the following way. Increasing the size of the aldehyde by using Isatin^[33]^ instead of nitrobenzaldehyde **1** with the aim to achieve stereoselectivity via “substrate engineering” led to conversions up to 87% and using the even bigger N-methyl-isatin conversion could be measured again but to no enhanced enantioselectivity was detected (Table S4). The lack of enantioselectivity of the catalyst was more deeply investigated using DFT methods. We employed ORCA^[34]^ to calculate the reaction pathway until Intermediate 2, where the chiral center is already formed^[35]^ (Figure 3, A) at a B3LYP level of theory using a dispersion correction^[36]^ and a CPCM model for water. As basis set ma-def2-SVP^[37]^ was employed.

**Figure 3.**
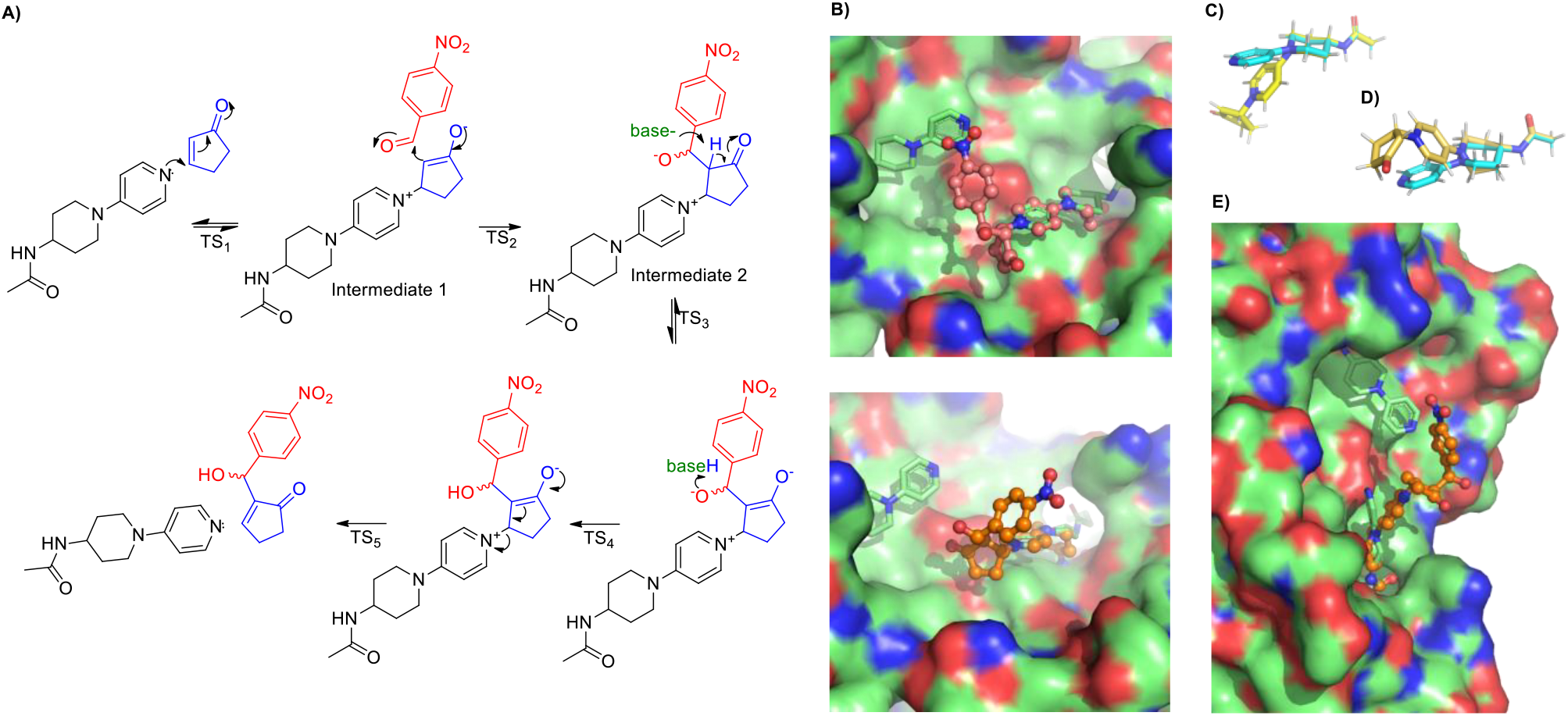
Explanation of stereoselective outcome of reaction using DFT calculated intermediates and the protein crystal structure of streptavidin S112I Q114A R121A L124Y. **A)** Proposed reaction mechanism of the Baylis-Hillman reaction **B)** Both possible enantiomers of Intermediate 2 overlaid on the pyridine ring of the catalyst **4**. Top: (*R*)-product (pink), Bottom: (*S*)-product (orange). **C)** DFT structure of N-(1-(pyridin-4-yl)piperidin-4-yl)acetamide (turquois) as catalyst and Intermediate 1 in chair conformation (yellow, right) and **D)** boat conformation (gold, left), respectively, overlaid on amide bond **E)** Intermediate 2 in twisted boat conformation overlaid on the amid bond from **4** in the crystal structure of the host protein. The cofactor is kinked and points towards the solvent.

By overlaying the positions of the pyridine ring of the catalyst of DFT-derived structures and the crystal structure, one could argue that the reasons for the missing enantioselectivity of the reaction should be clarified. As depicted in Figure 2, B the formation of both enantiomers is possible since both intermediates are not sterically hindered by surrounding amino acids although the intermediate leading to the (*S*) product (Figure 2 B, bottom) seems to undergo more favorable interactions with the protein (cyclopentenone **1** oxygen to Y124). Additionally, there is less spatial proximity of the cofactor **4** of chain B to -NO_2_ of 2 as compared to the intermediate leading to the (R)-product (Figure 2 B, top). This should at least lead to a moderate enantioselectivity, which we could not observe. If the proposed mechanism of action using DMAP as catalyst is considered in more detail, we notice that during the intermediate steps the positive charge of the pyridinium migrates to the amine-nitrogen atom (Figure S3), which leads to some consequences concerning cofactor **4**. Since cofactor **4** possesses at this position, in contrast to DMAP, a 6-membered ring the conformation of this piperidine ring has to change due to the need for accommodation of a sp^2^ hybridized amine nitrogen atom eventually bearing the positive charge. This conformational change propagates through **4** leading to a kinked molecule. Two possible conformations of the piperidine ring are possible – either the energetically favored chair conformation^[38]^ or the unfavored (twisted) boat conformation.

The energetically lower conformation of the piperidine possessing a chair conformation (Figure 3 C) is hindered by the surrounding protein. On the other hand, fits the twisted boat conformation of piperidine (Figure 3, D), around 3.8 kcal/mol higher in energy for intermediate 1, quite well into into the given protein environment. The conformational change places the reactive nitrogen at a very different spatial position as seen in almost all crystal structures (Figure 3, C-E). Since the protein crystallized in all cases, except the wildtype, under basic conditions with a neutral cofactor this conformation was never observed. Interestingly, the piperidine boat conformation of the catalyst is observed once in the solved crystal structures, namely in variant S112M Q114A R121A L124Y (PDB: 6T2Z, Figure S1). But only one of the two chains of the assymetric unit harbous **4** in this conformation. The second chain has electron density for **4** in a piperidine chair conformation. Surprisingly, this variant has the lowest activity of all active variants.

Since the catalyst is according to this theoretical considerations and calculations now pointing towards the solvent as depicted in Figure 3E, any chiral induction by the protein environment is lost, leading to achiral product formation.

The catalytic rate enhancement of protein with **4** over the rate of **4** is most presumably due to the hydrophobic pocket around the cofactor, increasing locally the concentration of the substrates and facilitating the formation of Intermediate 1.

In conclusion, a new artificial cofactor was developed, which utilizes a known organocatalyst and the biotin-streptavidin technology to successfully catalyze the Baylis-Hillman reaction. This protein-cofactor system yields a ~35-fold higher conversion compared to the bare artificial cofactor alone. The system was evolved via repeated cycles of mutagenesis, activity tests and structure determination. Hence, all protein design steps are rationalized. The reaction catalyzed by the artificial cofactor was further elucidated using DFT calculations to show major conformational changes of the catalyst, preventing the formation of chiral products. Here we demonstrate, how carefully considering the chemistry of a reaction, if fully understood, as well as deep knowledge of the (structural) properties of the target proteins are key to success in enzyme design.

## Acknowledgements

H.L. gratefully acknowledges financial support by the Austrian Science Fund (FWF, Erwin Schrödinger fellowship, J3994 B21). We thank Sooruban Shanmugaratnam for excellent technical assistance, all X-ray team members at the department for trips to Berlin, HZB (Berlin, Germany) for the allocation of synchrotron radiation beamtime at BESSY II as well as the Keylab HPC (University of Bayreuth) for computer time and the Central Analytics of the Organic Chemistry Department (University of Bayreuth) for HRMS measurements.

