## Supplement for "An Artificial Cofactor catalyzing the Baylis-Hillman Reaction using Designed Streptavidin as Protein Host"

### Synthesis

#### Chemicals

*N*-1-(pyridin-4-yl)piperidin-4-amine dihydrochloride and isatin was ordered at TCI. Biotin was from Roth. *N*-hydroxysuccinimide, *N,N'*-Dicyclohexylcarbodiimide, DMAP, cyclopentenone and 4-nitrobenzaldehyde were ordered from Sigma Aldrich.

#### Synthesis of ligand 1:

##### Step 1: Synthesis of Biotinyl-N-hydroxysuccinimide.

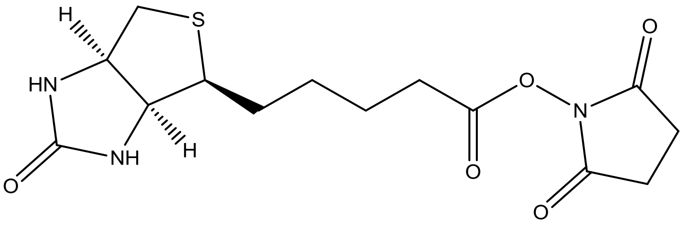

d-Biotin (0.25 g, 1.0 mmol) and *N*-hydroxysuccinimide (0.118 g, 1.0 mmol) were dissolved in hot DMF (15 mL). *N,N'*-Dicyclohexylcarbodiimide (0.220 g, 1.1 mmol) was added and the solution was stirred overnight at room temperature. A white precipitate was formed. The reaction mixture was filtered over celite. The product was crystallized from the filtrate by adding diethyl ether. After filtration and washing with diethyl ether a white solid was obtained (242 mg, 0.71 mmol, yield: 71%).

$^1$H NMR (500 MHz, DMSO) δ 6.44 (s, 1H), 6.38 (s, 1H), 4.35 – 4.27 (m, 1H), 4.19 – 4.11 (m, 1H), 3.16 – 3.06 (m, 1H), 2.87 – 2.76 (m, 5H), 2.68 (t, *J* = 7.4 Hz, 2H), 2.59 (d, *J* = 12.5 Hz, 1H), 1.72 – 1.56 (m, 3H), 1.56 – 1.35 (m, 3H). NMR according to literature. [1]

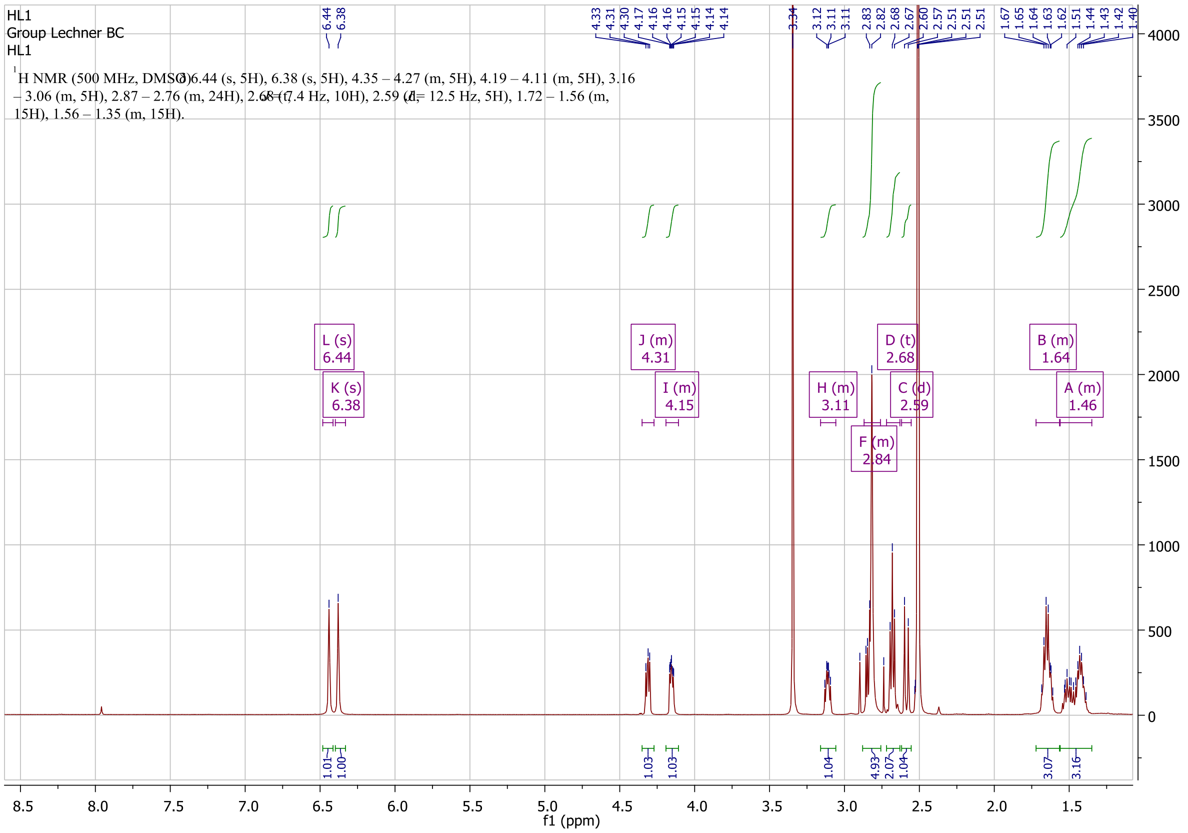

#### Step 2: Synthesis of Biotin-*N*-1-(pyridin-4-yl)piperidin-4-amine conjugate.

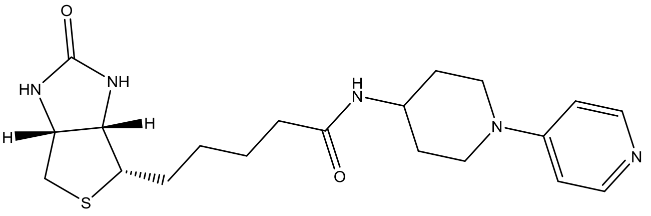

Biotinyl-N-hydroxysuccinimide (0.200 g, 0.59 mmol), triethyl amine (2.5 eq, 20 µL) and *N*-1-(pyridin-4-yl)piperidin-4-amine dihydrochloride (1.1 eq., 160 mg) were dissolved in DMF (25 mL) and the solution was stirred overnight at room temperature. The product was precipitated from the reaction with diethyl ether. A white precipitate was formed, which was filtered and washed with water. The solid was recrystallized from MeOH/diethyl ether within 3 days yielding the product (109 mg, 0.26 mmol, yield: 48%).

$^{1}$H NMR (500 MHz, DMSO) δ 8.13 (d, *J* = 6.3 Hz, 2H), 7.75 (d, *J* = 7.7 Hz, 1H), 6.82 (d, *J* = 6.4 Hz, 2H), 6.42 (s, 1H), 6.36 (s, 1H), 4.35 – 4.25 (m, 1H), 4.17 – 4.05 (m, 1H), 3.85 (d, *J* = 13.5 Hz, 4H), 3.12 – 3.06 (m, 1H), 2.96 (t, *J* = 11.4 Hz, 2H), 2.81 (dd, *J* = 12.4, 5.1 Hz, 1H), 2.57 (d, *J* = 12.4 Hz, 1H), 2.04 (t, *J* = 7.3 Hz, 2H), 1.85 – 1.68 (m, 2H), 1.52 (s, 2H), 1.52 – 1.04 (m, 7H).

$^{13}$C NMR (126 MHz, DMSO) δ 170.7, 162.1, 153.5, 149.0, 107.8, 60.4, 58.6, 54.8, 44.9, 44.0, 34.6, 29.9, 27.6, 27.4, 24.7.

HRMS (ESI): m/z calculated for $C_{20}H_{31}O_2N_5S$ [M+H]: 404.21147; found: 404.20978

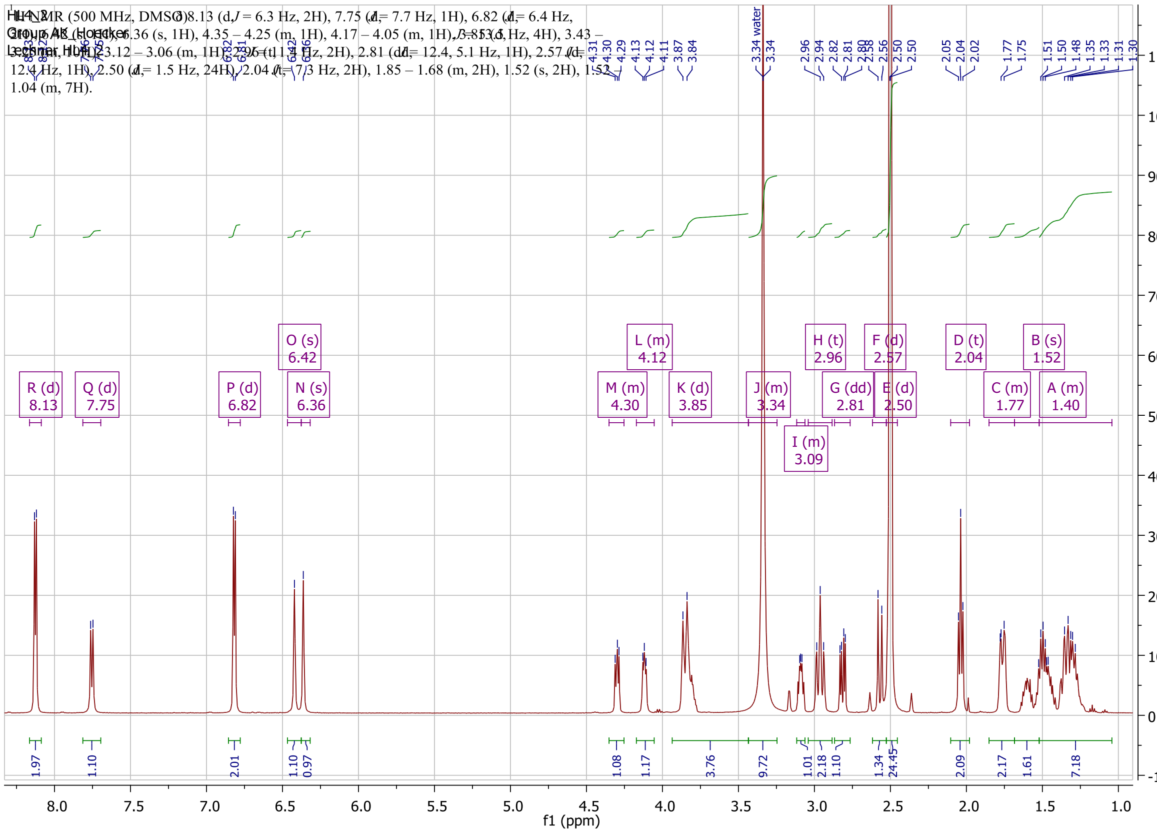
3.34 water
R (d) 8.13
Q (d) 7.75
P (d) 6.82
O (s) 6.42
N (s) 6.36
M (m) 4.30
L (m) 4.12
K (d) 3.85
J (m) 3.34
I (m) 3.09
H (t) 2.96
G (dd) 2.81
F (d) 2.57
E (d) 2.50
D (t) 2.04
C (m) 1.77
B (s) 1.52
A (m) 1.40
f1 (ppm)

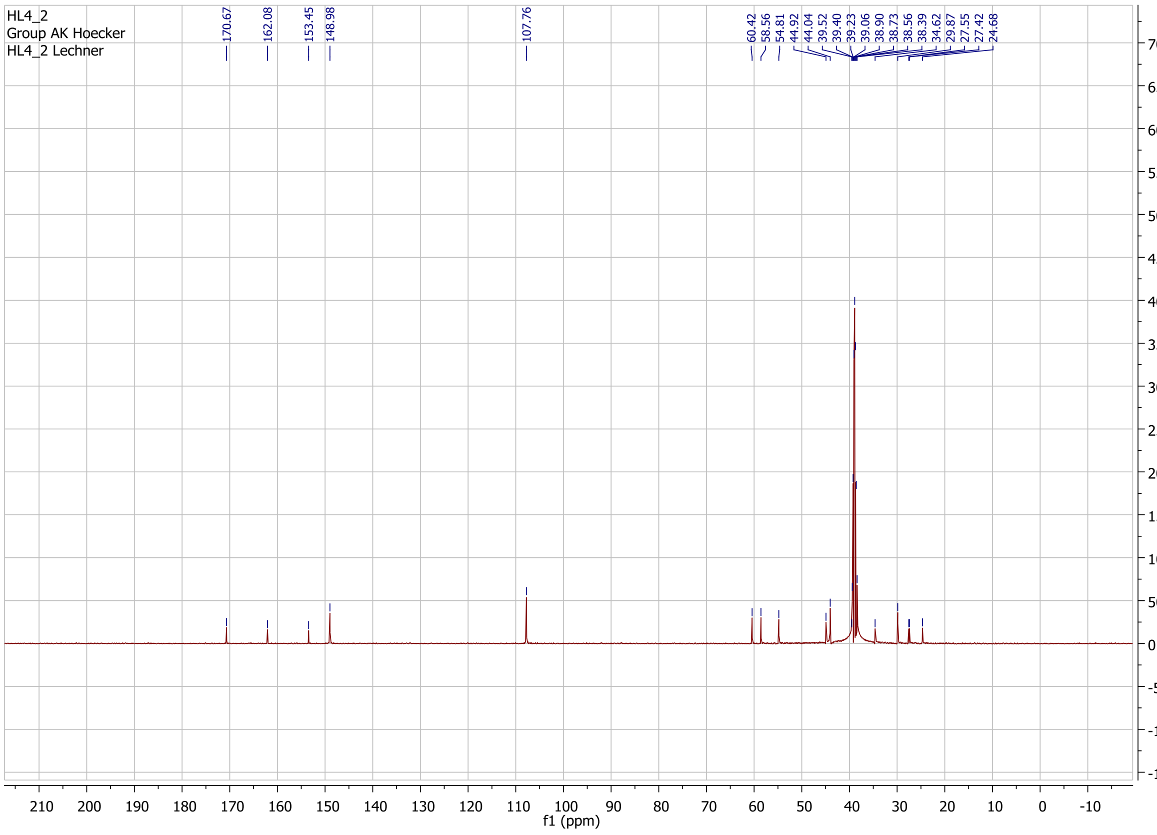
HL4_2
Group AK Hoecker
HL4_2 Lechner
f1 (ppm)

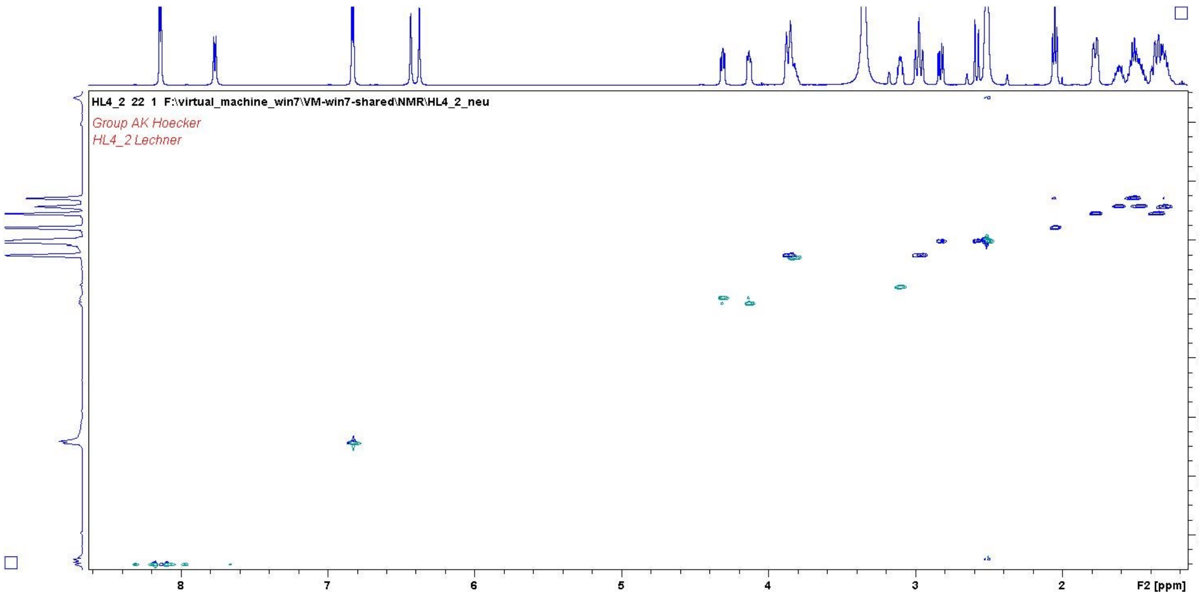

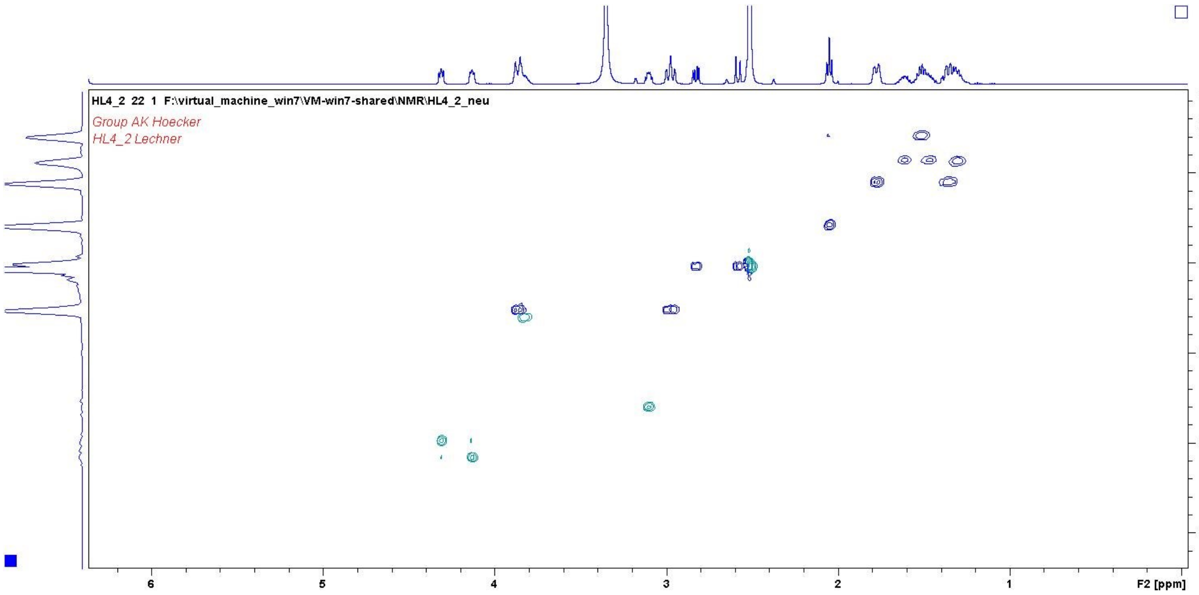

#### Synthesis of 2-(hydroxy(4-nitrophenyl)methyl)cyclopent-2-enone

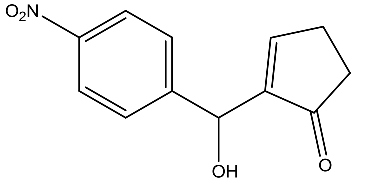

Procedure from [2]:
4-nitrobenzaldehyde (2 mmol, 302 mg) was dissolved in THF/$H_2O$ mixture (1.5 ml each), DMAP (20 mol%, 50 mg) and cyclopentenone (2.5 mmol, 205 µl) were added. The reaction mixture was extracted with ethyl acetate (3 x 5 ml). After adding a small amount of silica and evaporation of the solvent, the product was purified using a silica column. A gradient from hexanes/ethyl acetate of 99:1 to 7:3 yielded the product. According to NMR minor impurities were contained which could be removed by recrystallization from MeOH/acetonitrile (yellowish crystals, 155 mg, 0.66 mmol, yield 33%).

$^{1}H$ NMR (500 MHz, $CDCl_3$) δ 8.21 (d, *J* = 8.8 Hz, 2H), 7.58 (d, *J* = 8.6 Hz, 2H), 7.30 – 7.26 (m, 1H), 5.67 (d, *J* = 3.2 Hz, 1H), 3.56 (d, *J* = 4.4 Hz, 1H), 2.63 (m, 2H), 2.48 (m, 2H).

$^{13}C$ NMR (126 MHz, $CDCl_3$) δ 209.4, 159.8, 148.4, 147.5, 146.6, 127.1, 123.8, 69.2, 35.2, 26.9.

NMR according to Literature. [3]

HRMS (ESI): m/z calculated for $C_{12}H_{12}O_4N$ [M+H]: 234.07608; found: 234.07601

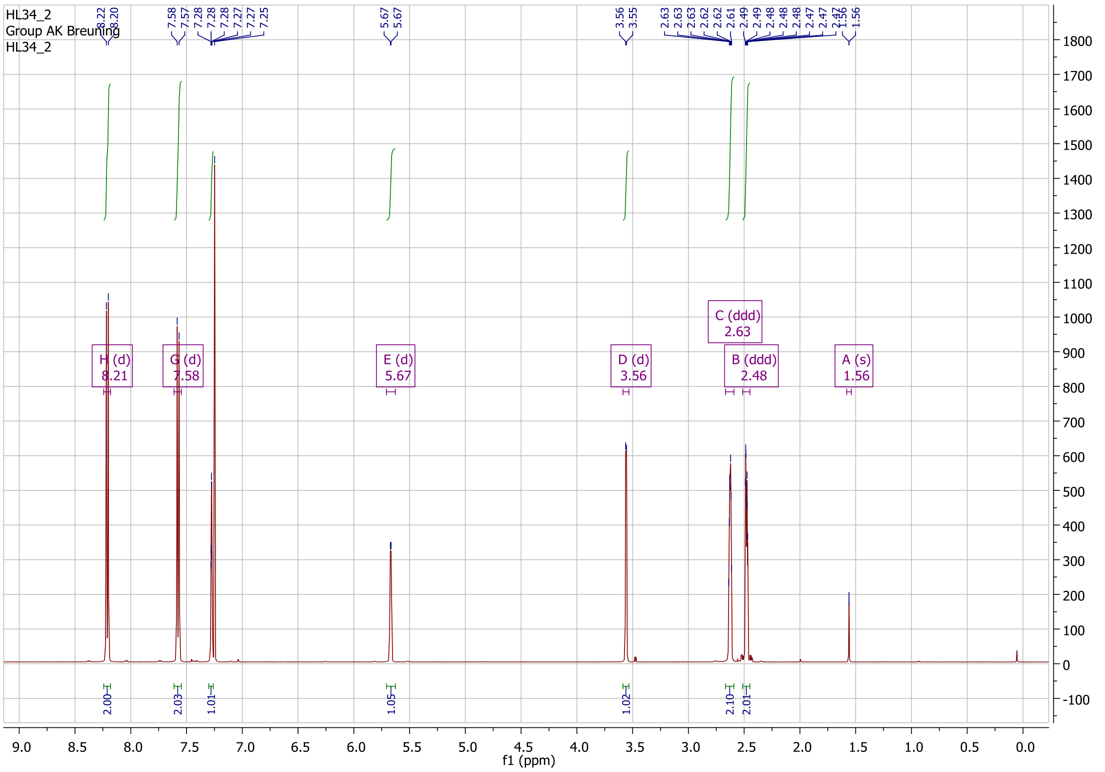

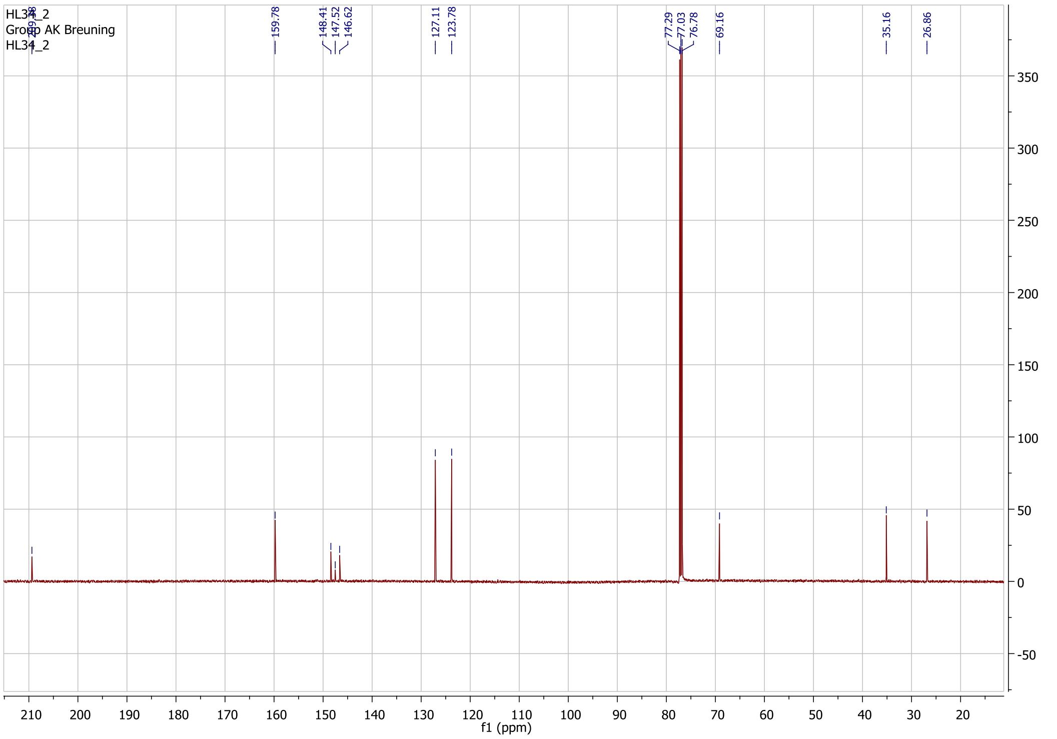

#### Synthesis of 3-hydroxy-3-(5-oxocyclopent-1-en-1-yl)indolin-2-one

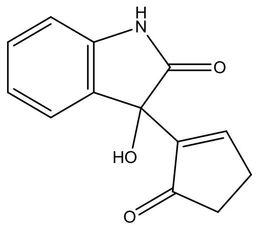

Procedure from [2]:
Isatin (2 mmol, 294 mg) was dissolved in THF/$H_2O$ mixture (1.5 ml each), DMAP (20 mol%, 50 mg) and cyclopentenone (2.5 mmol, 205 µl) were added. After stirring for 18 hours at 21°C the suspension was filtrated, and the solid residue was washed with hexanes (~20 ml). NMR of the product (yellowish, 150 mg, yield 32%) showed minor impurities. Hence the product was further purified using a silica column. A gradient from hexanes/ethyl acetate of 9:1 to 2:3 yielded finally pure product (white powder, 61 mg, 0.26 mmol, yield 14%).

$^1$H NMR (500 MHz, MeOD) δ 8.02 (t, *J* = 2.7 Hz, 1H), 7.23 (td, *J* = 7.7, 1.2 Hz, 1H), 7.13 (d, *J* = 7.8 Hz, 1H), 6.96 (td, *J* = 7.6, 0.8 Hz, 1H), 6.89 (d, *J* = 7.8 Hz, 1H), 2.76 – 2.69 (m, 2H), 2.45 – 2.29 (m, 2H).
$^{13}$C NMR (126 MHz, MeOD) δ 209.0, 180.0, 163.2, 163.2, 146.9, 144.0, 132.0, 131.1, 125.5, 123.7, 111.5, 76.1, 36.5, 28.0.
NMR according to literature. [4]

HRMS (ESI): m/z calculated for $C_{13}H_{11}O_3N_1Na$ [M+Na]: 252.06311; found: 252.06213

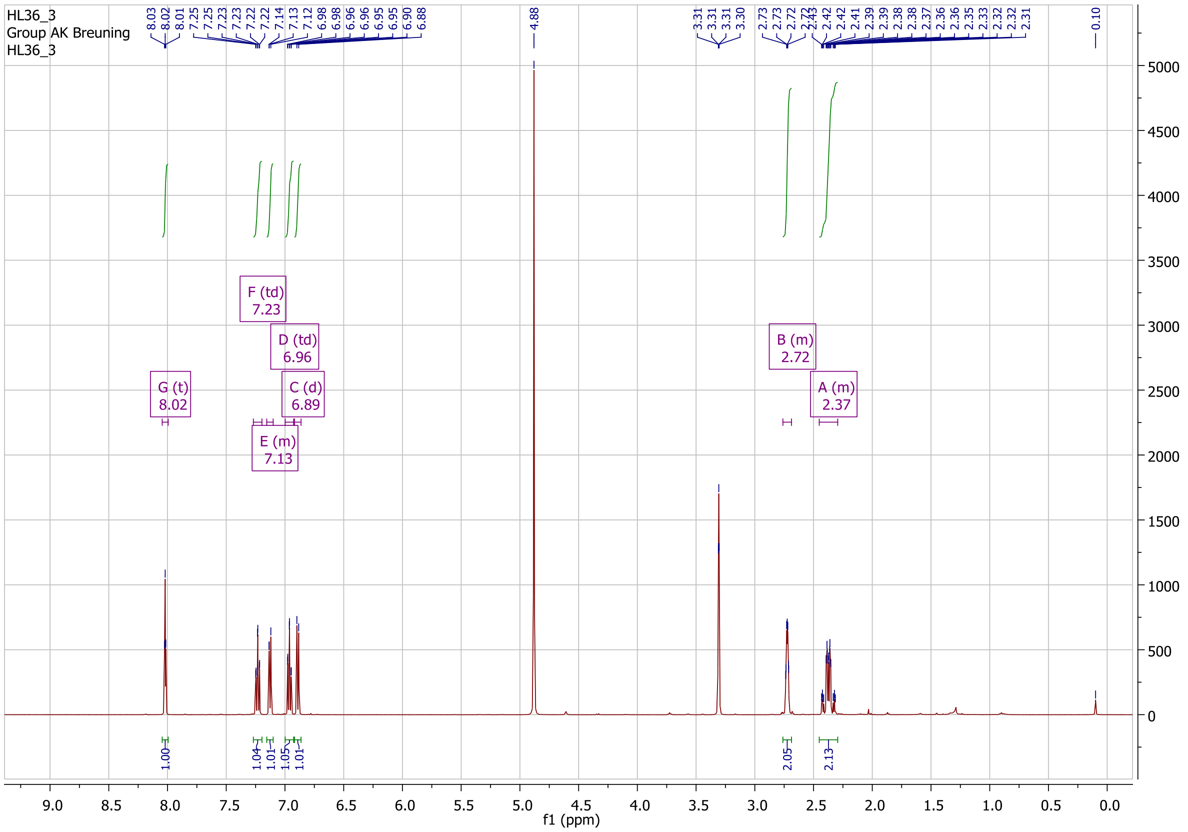

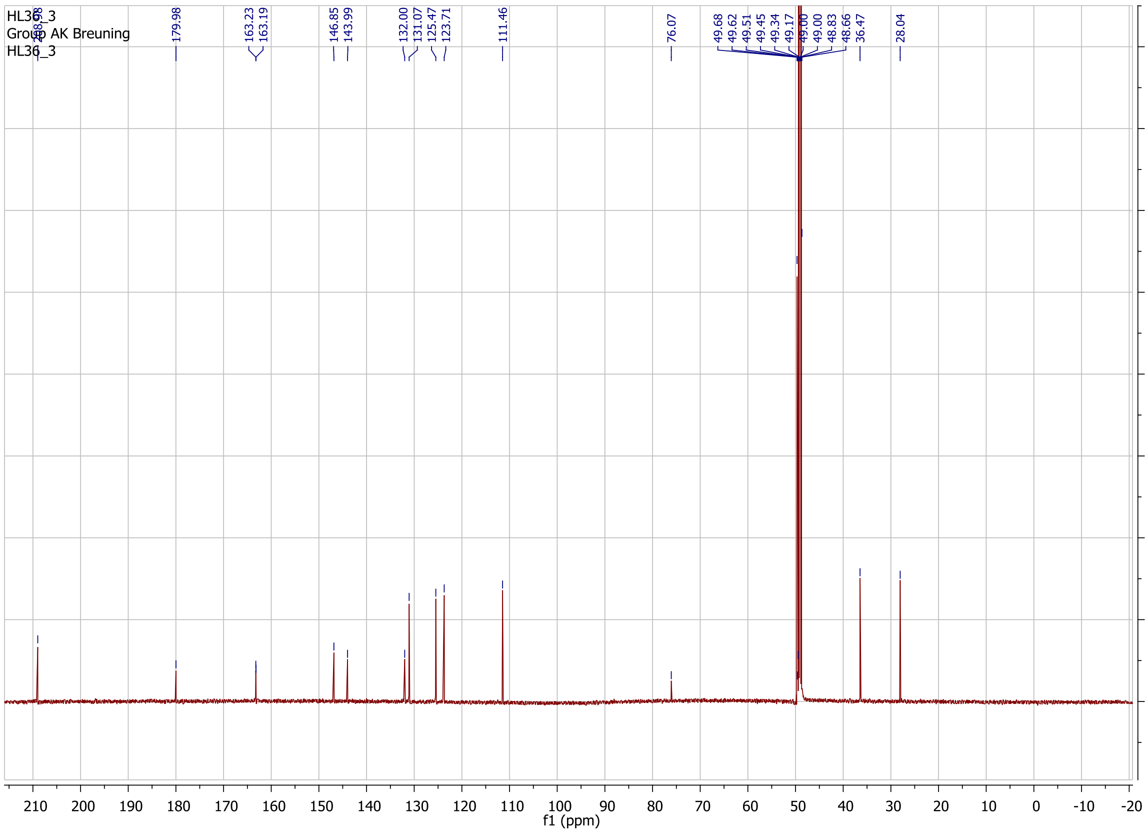

HL36_3
Group AK Breuning
HL36_3
208.98
179.98
163.23
163.19
146.85
143.99
132.00
131.07
125.47
123.71
111.46
76.07
49.68
49.62
49.51
49.45
49.34
49.17
49.00
49.00
48.83
48.66
36.47
28.04
f1 (ppm)

#### Streptavidin-sequence

Streptavidin (PDB: 5F2B)[5] with following sequence was ordered as synthetic gene as wt-streptavidin.

MASMTGGQQMGRDEAGITGTWYNQLGSTFIVTAGADGALTGTYESAAGKAESRYVLTGRYDSAPATDGSGTAL
GWTVAWKNNYRNAHSATTWSGQYVGGAEARINTQWLLTSGQTEANGWRSTLVGHDTFTKVKPSAASIDAAKK
AGVNNGNPLDAVQQ

catatggctagtatgacaggcggacaacaaatgggtcgtgacgaagcaggcatcaccggcacttggtacaatcaacttggttcaactttcatcgt
cacagcgggtgccgacggtgcgcttacagggacttatgaatcagcagccgggaaggccgaaagtcgttatgttttgacaggccgctacgactcc
gcgccggcaactgacggtagcggaacagcactgggctggacagttgcttggaaaaataattatcgtaacgctcattctgctacaacctggagcg
gtcagtacgtcggcggtgctgaggctcgtatcaacactcaatggttgttaaccagcgggcaaactgaagctaatggatggcgcagcactttagtc
ggacacgacaccttcacgaaagttaaaccttcggctgcgagcatcgatgccgcgaaaaaggctggtgttaataatggcaatcctcttgatgcagt
gcagcagtaaggatcc

#### X-ray crystallography

##### Structure determination

Data collection was done at BESSY II beamline MX14.1/2 (operated by the Helmholtz Zentrum Berlin, Germany) and a Pilatus 6M/2M detector (Dectris, Baden, Switzerland). Indexing, scaling and merging of diffraction data was performed with XDSapp[6] based on XDS.[7] For structure solving the following packages as included in Phenix[8] were used: Structures were solved by molecular replacement with PHASER[9] using as a search model for the wt-streptavidin an existing streptavidin structure (PDB-Code 2qcb). Subsequent molecular replacement calculations for the other structures were done using the derived wt-streptavidin structure. Refinement was performed using phenix.refine.[10] For the wt-streptavidin, variants S112A Q114A R121A and S112F Q114A R121A L124Y anisotropic refinement was advantageous. Manual model building was done using coot.[11]
The ligand was optimized and a cif file created using GAMESS[12] and phenix.eLBOW. For graphical representation Pymol (Schrodinger LCC) was used.

##### Wt-streptavidin with ligand 4 (soaked) (PDB: 6T1E)

Streptavidin (16.6 mg/ml, 1 µl, in HEPES 10 mM, pH 8) was added to the same amount of crystallization buffer in a hanging drop vapor-diffusion approach. The conditions of pdb:2bc3 [13] were reproduced, but also the condition slightly varied.

After one day crystals were observed in all conditions. On day 2, crystals were soaked with ligand **1** (stock 10 mM in DMSO). The ligand solution (1 µl) was mixed with crystallization buffer ($(NH_4)_2SO_4$ 1.8 M, $NaCH_3COO$ 0.1 M, pH 4.6, 9 µl) and crystals were transferred into this drop. After 1 hour of equilibration against the crystallization buffer the crystals were transferred in a drop containing $(NH_4)_2SO_4$ 1.8 M, $NaCH_3COO$ 0.1 M, pH 4.6 and glycerol (25 %) and flash-frozen in liquid nitrogen.

Soaking in same buffer, but pH 8.0 and 7.0 lead to dissolving of the crystal. With pH 6.0 and a soaking time of 20 min the crystal was present but was very fragile and could not be mounted.

**Table S1**. Data collection and refinement statistics of streptavidin with biotinylated 4-(piperidin-1-yl)pyridine. (PDB: 6T1E).

| | |
|---|---|
| **Wavelength** | 0.918400 |
| **Resolution range** | 28.76 - 1.30 (1.35 - 1.30) |
| **Space group** | I 41 2 2 |
| **Unit cell** | 57.51 57.51 174.66 90 90 90 |
| **Total reflections** | 525359 (49510) |
| **Unique reflections** | 36587 (3573) |
| **Multiplicity** | 14.4 (13.9) |
| **Completeness (%)** | 99.9 (99.9) |
| **Mean I/sigma(I)** | 18.71 (1.03) |
| **Wilson B-factor** | 17.00 |
| **R-merge** | 0.088 (2.433) |
| **R-meas** | 0.091 (2.527) |
| **R-pim** | 0.024 (0.669) |
| **CC1/2** | 1.000 (0.398) |
| **CC*** | 1.000 (0.755) |
| **Reflections used in refinement** | 36575 (3571) |
| **Reflections used for R-free** | 1829 (179) |
| **R-work** | 0.129 (0.302) |
| **R-free** | 0.158 (0.290) |
| **CC(work)** | 0.976 (0.730) |
| **CC(free)** | 0.966 (0.718) |
| **Number of non-hydrogen atoms** | 1185 |
| **macromolecules** | 1018 |
| **ligands** | 28 |
| **solvent** | 116 |
| **Protein residues** | 125 |
| **RMS(bonds)** | 0.016 |
| **RMS(angles)** | 1.36 |
| **Ramachandran favored (%)** | 98.37 |
| **Ramachandran allowed (%)** | 1.36 |
| **Ramachandran outliers (%)** | 0.00 |
| **Rotamer outliers (%)** | 0.00 |
| **Clash score** | 4.87 |
| **Average B-factor** | 24.68 |
| **macromolecules** | 22.72 |
| **ligands** | 19.70 |
| **solvent** | 38.43 |

Statistics for the highest-resolution shell are shown in parentheses.

#### Streptavidin variants with ligand 4 (cocrystallized)

Ligand **1** (100µl, stock 25mM in DMSO) was added to protein (500µl, streptavidin S112A Q114T R121A: 18.4 mg/ml, streptavidin S112A R121A: 20.4 mg/ml, streptavidin S112A Q114A R121A: 20.0 mg/ml, streptavidin S112A Q114A R121A L124Y 16 mg/ml). After incubation (1hour) at RT a NAP-5 column was used to purify and change the buffer to HEPES (10 mM, pH 8.0). The amount of liquid was reduced to 500µl again.

Screening in a 96well, 3 sitting drop well format using the JCSG Core [14] I-IV screen (0.4µl protein+0.4µl buffer) yielded in crystals in the above listed conditions. If necessary, the crystal was transferred to a drop of reservoir solution supplemented with 25% glycerol for 1 min. Crystals were flash-frozen in liquid nitrogen.

- **PDB 6T1G** (S112A Q114T R121A): Na-citrate 0.1 M pH 5.5, 40% PEG 600
- **PDB 6T1K** (S112A R121A): CHES 0.1 M pH 9.5 30% PEG 3000
- **PDB 6T2L** (S112A Q114A R121A): Na-acetate 0.17 M, TRIS 0.085 M pH 8.5, PEG 4000 25.5%, glycerol 15%
- **PDB 6T2Y** (S112A Q114A R121A L124Y): di-Sodium hydrogen phosphate 0.2M, PEG 3350 20%; cryo: glycerol 25%

S112A Q114T R121A

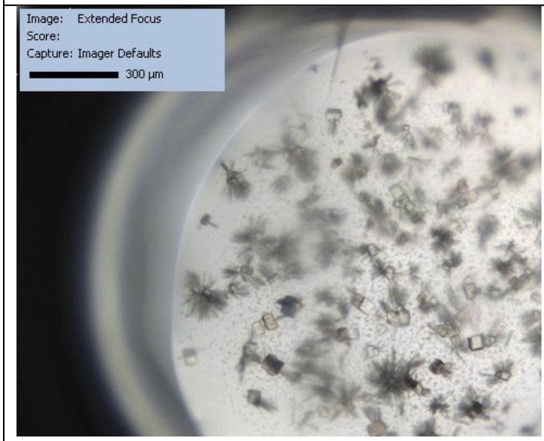

S112A R121A

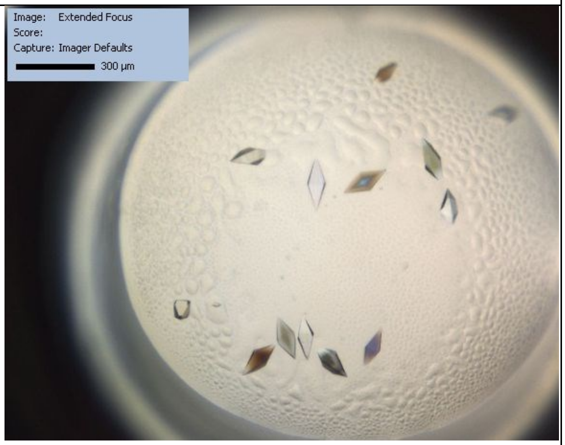

S112A Q114A R121A

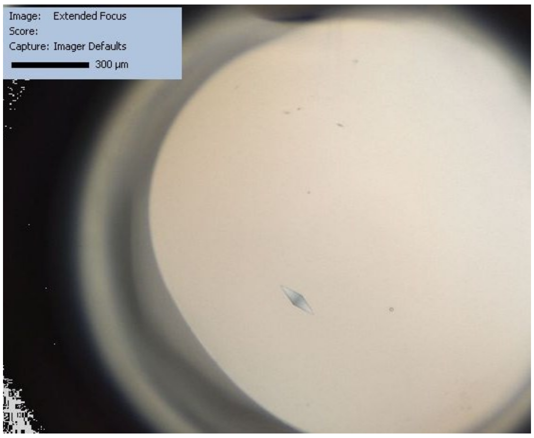

S112A Q114A R121A L124Y

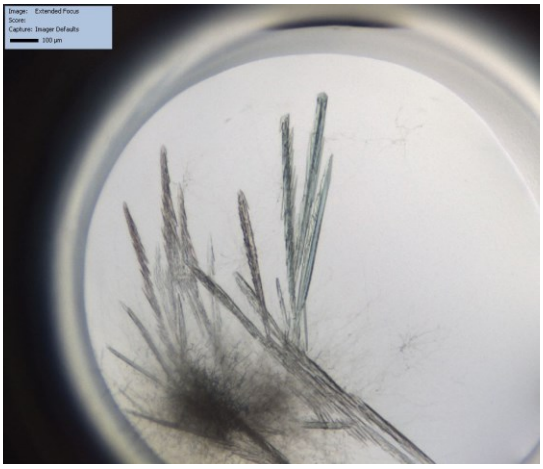

**Table S2.** Data collection and refinement statistics of streptavidin variants with biotinylated 4-(piperidin-1-yl)pyridine.

| | 6T1G (S112A Q114T R121A) | 6T1K (S112A R121A) | 6T2L (S112A Q114A R121A) | 6T2Y (S112A Q114A R121A L124Y) |
|---|---|---|---|---|
| **Wavelength** | 0.918400 | 0.918400 | 0.918400 | 0.918400 |
| **Resolution range** | 46.19 - 1.90 (1.97 - 1.90) | 43.75 - 1.20 (1.24 - 1.20) | 43.72 - 1.00 (1.04 - 1.00) | 47.54 - 1.80 (1.86 - 1.80) |
| **Space group** | P 43 2 2 | I 41 2 2 | I 41 2 2 | C 2 2 21 |
| **Unit cell** | 55.23 55.23 168.48 90 90 90 | 57.28 57.28 175.02 90 90 90 | 57.22 57.22 174.89 90 90 90 | 57.52 84.45 98.67 90 90 90 |
| **Total reflections** | 301408 (29252) | 493324 (45198) | 662469 (61368) | 167832 (16920) |
| **Unique reflections** | 21439 (2055) | 46054 (4461) | 78168 (7610) | 41969 (4149) |
| **Multiplicity** | 14.1 (14.2) | 10.7 (10.1) | 8.5 (8.1) | 7.6 (7.9) |
| **Completeness (%)** | 99.4 (98.4) | 99.8 (98.9) | 99.5 (98.3) | 97.7 (97.3) |
| **Mean I/sigma(I)** | 10.16 (0.68) | 10.47 (0.77) | 9.49 (0.70) | 4.80 (0.58) |
| **Wilson B-factor** | 31.80 | 14.78 | 9.19 | 31.77 |
| **R-merge** | 0.228 (4.354) | 0.127 (2.784) | 0.121 (2.280) | 0.2678 (2.650) |
| **R-meas** | 0.236 (4.515) | 0.134 (2.932) | 0.129 (2.436) | 0.2877 (2.835) |
| **R-pim** | 0.063 (1.179) | 0.041 (0.911) | 0.043 (0.842) | 0.104 (1.000) |
| **CC1/2** | 0.999 (0.519) | 0.999 (0.301) | 0.998 (0.356) | 0.985 (0.229) |
| **CC*** | 1.000 (0.827) | 1.000 (0.680) | 1.000 (0.725) | 0.996 (0.611) |
| **Reflections used in refinement** | 21392 (2052) | 46043 (4459) | 78155 (7609) | 41969 (4149) |
| **Reflections used for R-free** | 1069 (103) | 2100 (203) | 2101 (205) | 1108 (108) |
| **R-work** | 0.204 (0.352) | 0.157 (0.323) | 0.128 (0.441) | 0.193 (0.360) |
| **R-free** | 0.250 (0.4128) | 0.171 (0.323) | 0.138 (0.430) | 0.240 (0.338) |
| **CC(work)** | 0.953 (0.752) | 0.974 (0.630) | 0.982 (0.716) | 0.964 (0.545) |
| **CC(free)** | 0.921 (0.659) | 0.966 (0.612) | 0.971 (0.717) | 0.935 (0.463) |
| **Number of non-hydrogen atoms** | 2140 | 1247 | 1233 | 2052 |
| **macromolecules** | 1920 | 1050 | 1009 | 1840 |
| **ligands** | 56 | 28 | 28 | 56 |
| **solvent** | 160 | 157 | 186 | 142 |
| **Protein residues** | 256 | 123 | 121 | 244 |
| **RMS(bonds)** | 0.014 | 0.008 | 0.008 | 0.004 |
| **RMS(angles)** | 1.73 | 1.06 | 1.05 | 0.76 |
| **Ramachandran favored (%)** | 97.22 | 97.46 | 97.41 | 97.50 |
| **Ramachandran allowed (%)** | 2.38 | 2.54 | 2.59 | 2.50 |
| **Ramachandran outliers (%)** | 0.40 | 0.00 | 0.00 | 0.00 |
| **Rotamer outliers (%)** | 0.00 | 0.00 | 0.00 | 0.00 |
| **Clash score** | 5.03 | 6.13 | 4.41 | 3.02 |
| **Average B-factor** | 43.47 | 22.63 | 17.73 | 43.16 |
| **macromolecules** | 43.62 | 20.69 | 14.19 | 43.08 |
| **ligands** | 34.82 | 20.62 | 10.46 | 35.86 |
| **solvent** | 44.06 | 34.46 | 36.58 | 46.61 |
| **Number of TLS groups** | 13 | 1 | 1 | 13 |

Statistics for the highest-resolution shell are shown in parentheses.

Ligand **1** (100µl, stock 25mM in DMSO) was added to protein (500µl, streptavidin S112M Q114A R121A: 15 mg/ml, streptavidin Q114A R121A: 20 mg/ml, streptavidin S112F Q114A R121A L124Y: 18.0 mg/ml, streptavidin S112I Q114A R121A L124Y: 30mg/ml). After incubation (1hour) at RT a NAP-5 column was used to purify and change the buffer to HEPES (10 mM, pH 8.0). The amount of liquid was reduced to 500µl again.

Screening in a 96well, 3 sitting drop well format using JCSG Core [14] I-IV screen (0.4µl protein+0.4µl buffer) yielded in crystals in the above listed conditions. If necessary, the crystal was transferred to a drop of reservoir solution supplemented with 25% glycerol for 1 min. Crystals were flash-frozen in liquid nitrogen.

- **PDB 6T2Z** (S112M Q114A R121A): 1.6M $Na_2HPO_4$ 0.4M $K_2HPO_4$ 0.1M Sodium citrate phosphate pH 4.2
- **PDB 6T30** (Q114A R121A): 0.2M di-Sodium hydrogen phosphate 20% PEG 3350, cryo: glycerol 25%
- **PDB 6T31** (S112F Q114A R121A L124Y): CHES 0.1 M pH 9.5 30% PEG 3000
- **PDB 6T32** (S112I Q114A R121A L124Y): CHES 0.1 M pH 9.5 30% PEG 3000

<table>
<tr><td>S112M Q114A R121A</td><td>Q114A R121A</td></tr>
<tr><td>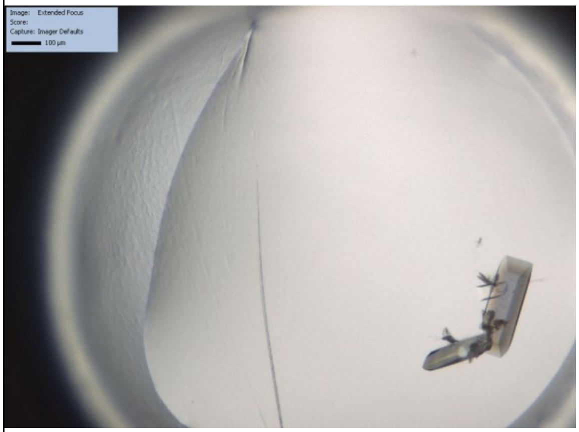</td><td>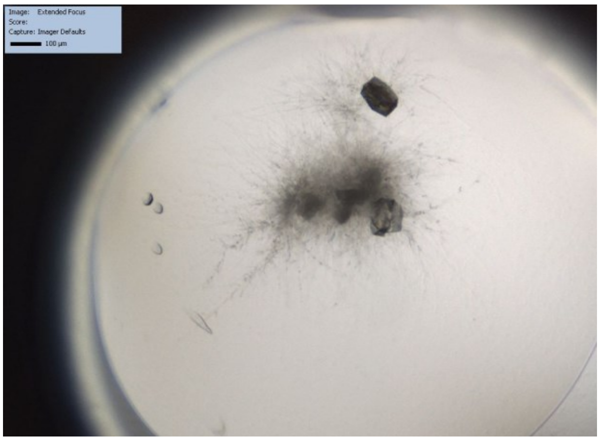</td></tr>
<tr><td>S112F Q114A R121A L124Y</td><td>S112I Q114A R121A L124Y</td></tr>
<tr><td>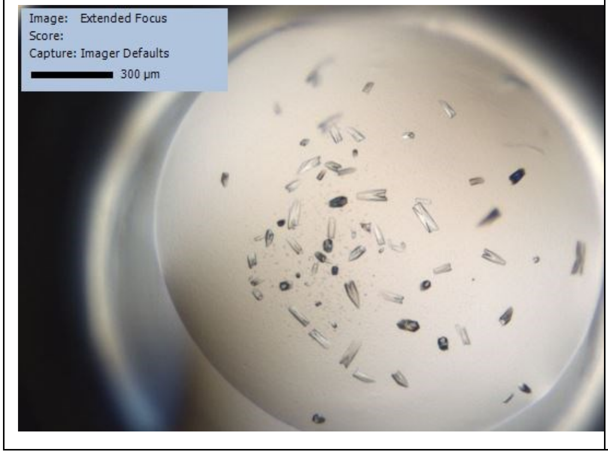
</td><td>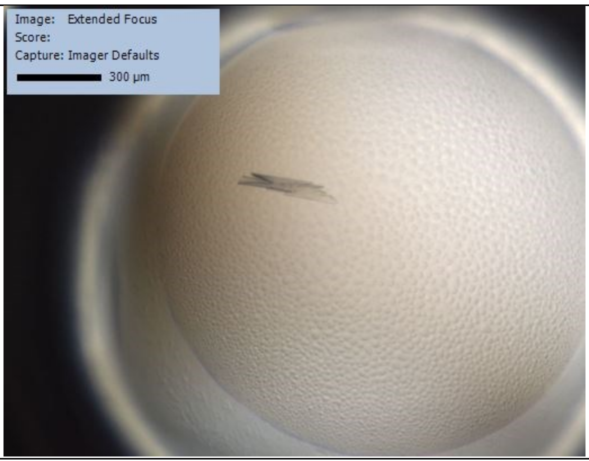
</td></tr>
</table>

**Table S3.** Data collection and refinement statistics of streptavidin variants with biotinylated 4-(piperidin-1-yl)pyridine.

| | 6T2Z (S112M Q114A R121A) | 6T30 (Q114A R121A) | 6T31 (S112F Q114A R121A L124Y) | 6T32 (S112I Q114A R121A L124Y) |
|---|---|---|---|---|
| Wavelength | 0.918400 | 0.918400 | 0.918400 | 0.918400 |
| Resolution range | 42.81 - 1.35 (1.34 - 1.35) | 47.75 - 1.80 (1.87 - 1.80) | 43.02 - 1.35 (1.40 - 1.35) | 42.54 - 1.75 (1.81 - 1.75) |
| Space group | C 2 2 21 | P 32 2 1 | C 2 2 21 | C 2 2 21 |
| Unit cell | 57.38 84.54 98.99 90 90 90 | 55.13 55.13 131.21 90 90 120 | 57.38 85.36 100.38 90 90 90 | 57.33 85.09 99.44 90 90 90 |
| Total reflections | 345405 (29194) | 238169 (24033) | 401552 (39888) | 118546 (11181) |
| Unique reflections | 52409 (4879) | 21936 (2112) | 54426 (5353) | 24739 (2318) |
| Multiplicity | 6.6 (6.0) | 10.9 (11.3) | 7.4 (7.5) | 4.8 (4.8) |
| Completeness (%) | 98.66 (93.06) | 98.89 (97.60) | 99.92 (99.61) | 98.47 (94.23) |
| Mean I/sigma(I) | 9.27 (0.71) | 17.36 (0.61) | 14.80 (1.45) | 10.05 (1.25) |
| Wilson B-factor | 22.96 | 38.93 | 18.61 | 27.85 |
| R-merge | 0.087 (1.689) | 0.078 (4.035) | 0.063 (1.114) | 0.084 (1.058) |
| R-meas | 0.094 (1.850) | 0.081 (4.223) | 0.068 (1.198) | 0.094 (1.184) |
| R-pim | 0.036 (0.741) | 0.025 (1.236) | 0.025 (0.435) | 0.042 (0.524) |
| CC1/2 | 0.996 (0.349) | 0.999 (0.366) | 0.999 (0.609) | 0.998 (0.556) |
| CC* | 0.999 (0.719) | 1.000 (0.732) | 1.000 (0.870) | 0.999 (0.845) |
| Reflections used in refinement | 52405 (4879) | 21916 (2111) | 54424 (5353) | 24731 (2318) |
| Reflections used for R-free | 2098 (195) | 1094 (106) | 2100 (206) | 1237 (116) |
| R-work | 0.164 (0.400) | 0.186 (0.373) | 0.138 (0.288) | 0.161 (0.338) |
| R-free | 0.182 (0.395) | 0.230 (0.400) | 0.165 (0.326) | 0.191 (0.376) |
| CC(work) | 0.967 (0.631) | 0.960 (0.646) | 0.975 (0.832) | 0.972 (0.788) |
| CC(free) | 0.963 (0.639) | 0.969 (0.688) | 0.958 (0.748) | 0.953 (0.814) |
| Number of non-hydrogen atoms | 2217 | 2035 | 2273 | 2105 |
| macromolecules | 1868 | 1845 | 1944 | 1852 |
| ligands | 56 | 56 | 56 | 56 |
| solvent | 293 | 116 | 273 | 197 |
| Protein residues | 243 | 248 | 246 | 242 |
| RMS(bonds) | 0.003 | 0.014 | 0.008 | 0.012 |
| RMS(angles) | 0.73 | 1.21 | 0.97 | 1.11 |
| Ramachandran favored (%) | 97.07 | 96.72 | 97.52 | 97.06 |
| Ramachandran allowed (%) | 2.93 | 3.28 | 2.48 | 2.52 |
| Ramachandran outliers (%) | 0.00 | 0.00 | 0.00 | 0.42 |
| Rotamer outliers (%) | 0.55 | 0.00 | 0.00 | 0.56 |
| Clash score | 1.91 | 4.66 | 3.92 | 1.37 |
| Average B-factor | 34.73 | 50.28 | 27.79 | 33.23 |
| macromolecules | 33.05 | 49.76 | 26.38 | 32.22 |
| ligands | 37.72 | 47.49 | 20.40 | 30.20 |
| solvent | 44.89 | 56.38 | 39.35 | 43.58 |
| Number of TLS groups | 1 | 1 | 11 | 1 |

Statistics for the highest-resolution shell are shown in parentheses.

#### Composite Omit Maps of ligands

Native (PDB: 6T1E)

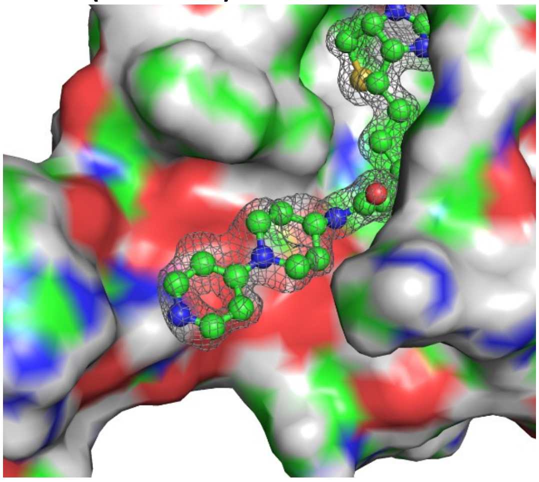

S112A Q114T R121A (PDB: 6T1G)

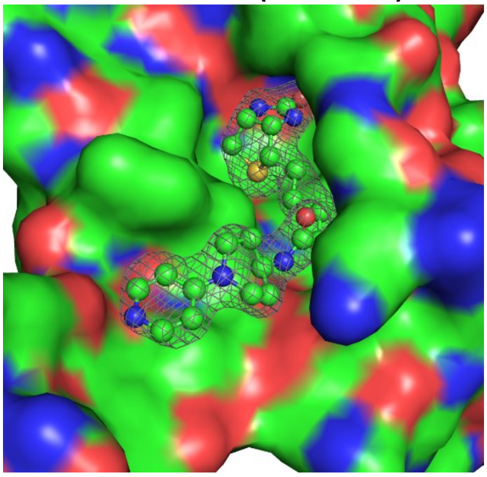

S112A R121A (PDB: 6T1K)

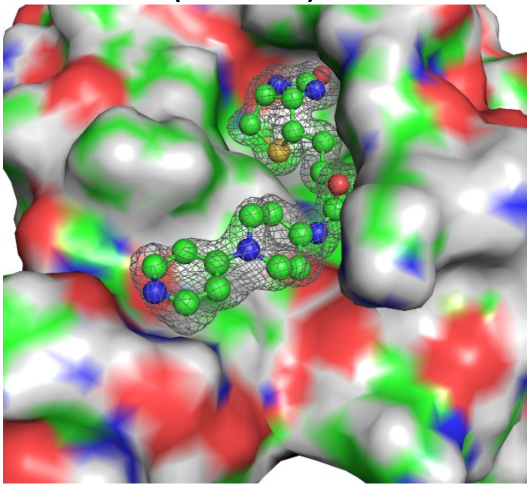

S112A Q114A R121A (PDB: 6T2L)

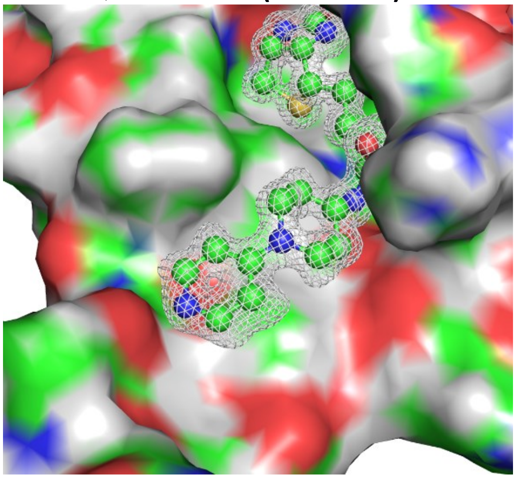

S112A Q114A R121A L124Y (PDB: 6T2Y)

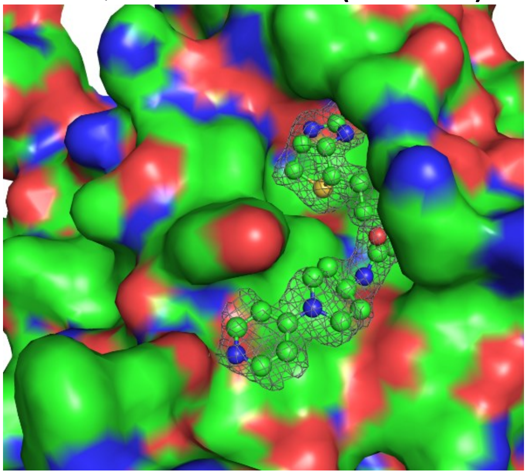

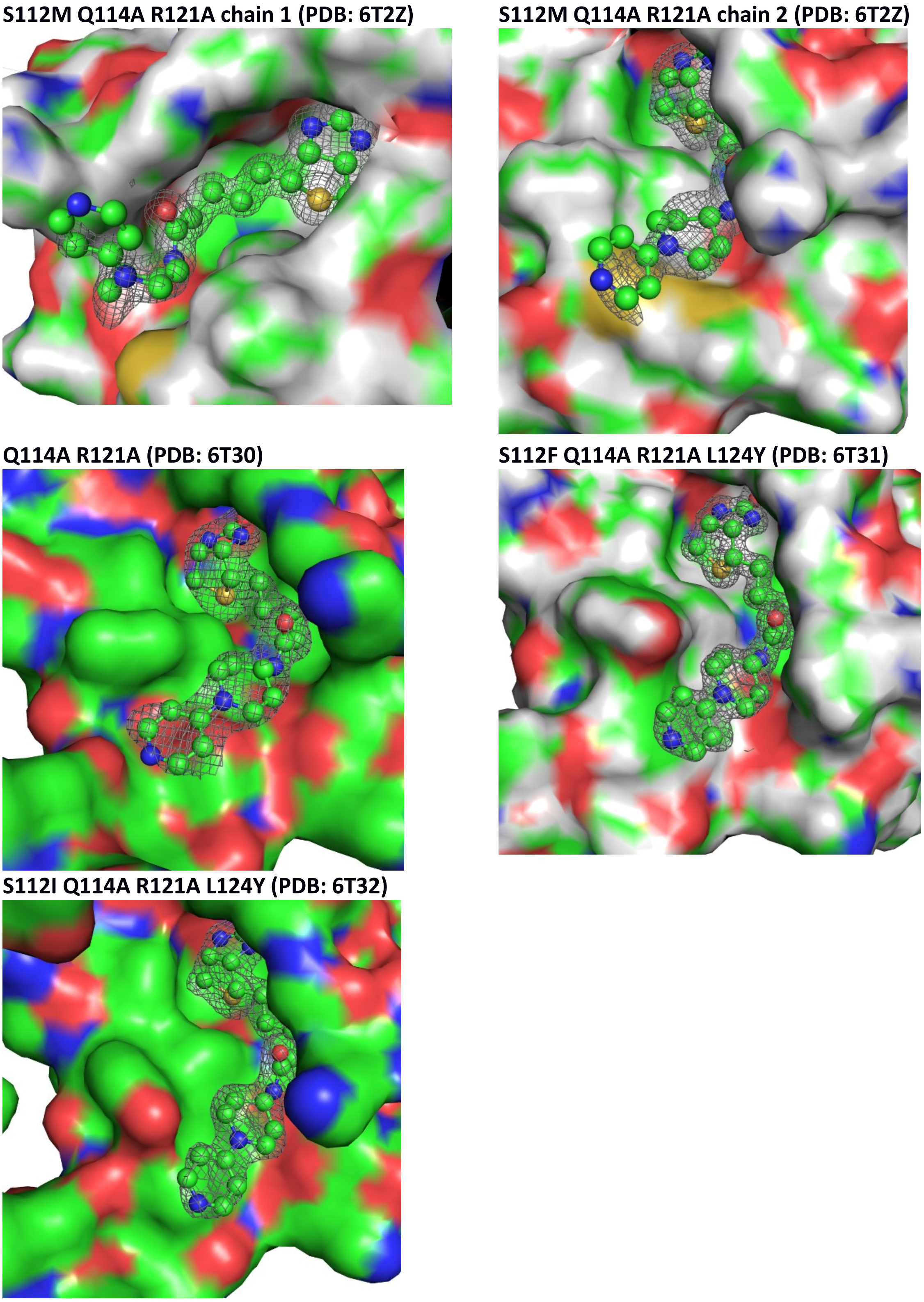

**Figure S1.** A simulated annealing composite omit map at sigma 1.0 level of the biotin-dmap ligand. Maps were created using Phenix[8] with 2mFo-DFc maps.

#### Histidine occurrence

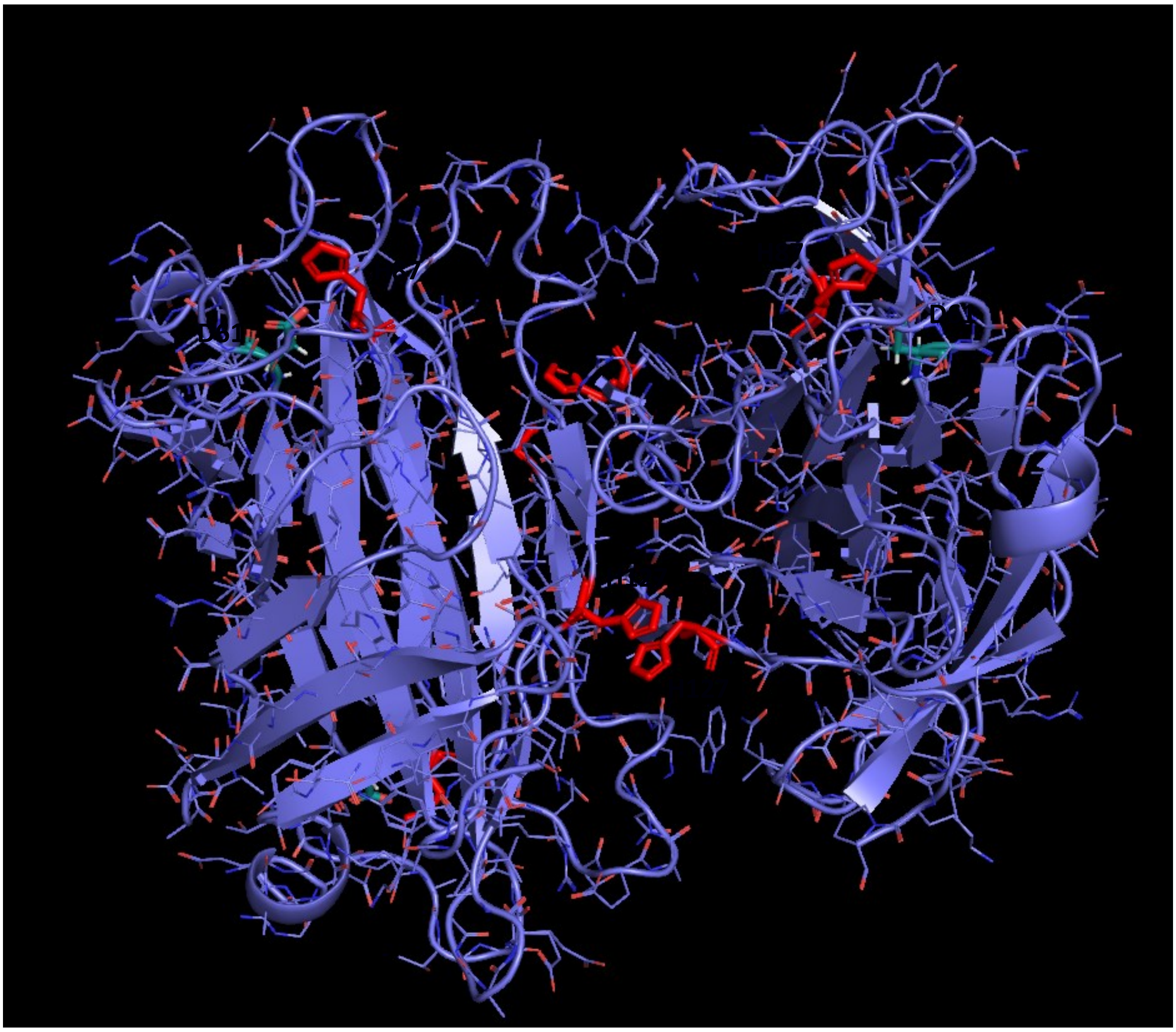

**Figure S2**. Position of histidine residues in wt-streptavidin.

#### Isomerization of DMAP

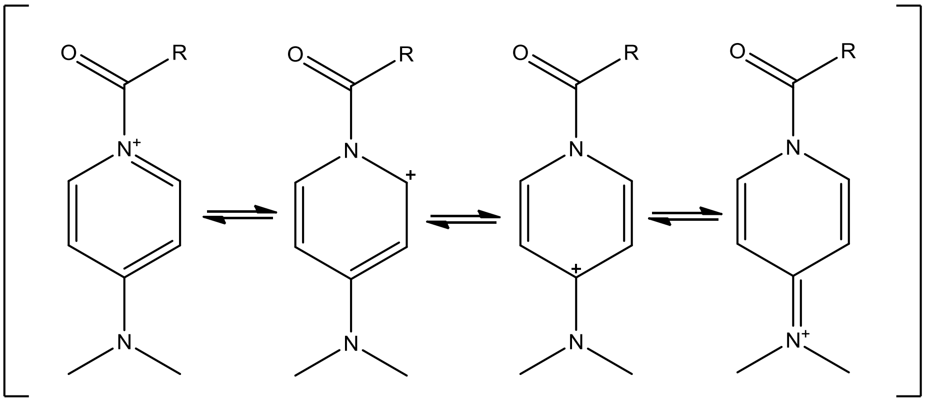

**Figure S3.** Cation isomeriszation on DMAP-substrate intermediate.

### DFT-calculations

All calculations were carried out using ORCA 4.1.0. [15,16] Transition sate search was done using the nudged elastic band method. Free energy corrections were determined using B3LYP with dispersion correction[17] and the ma-def2-SVP[18] as basis set using a the CPCM implicit solvation model for the solvent (water). Vibrational frequencies assuming a standard state of 1 atm and 298.15 K were calculated. Single point energy calculations were performed on optimized geometries with B3LYP/DB3J/def2-TZVP in conjunction with the CPCM model. The free energies displayed in Figure S4 were determined by adding zero-point energy and thermal correction determined using B3LYP/DB3J/def2-SVP/CPCM to electronic energies computed at the B3LYP/DB3J/def2-TZVP/CPCM level of theory. The Gibbs free energies were calculated at B3LYP/DB3J/def2-SVP/CPCM level of theory. They are in accordance with calculations carried out by Plata and Singleton[19], but as they conclude in their work it is rather difficult to obtain exact energies and geometries using these methods. For our case, the geometries should be reasonable enough to explain the outcome of our experiments.

We were interested in the steps until I2 (Figure 3A) , since during these steps the chirality is determined. For simplification was catalyst **4** truncated and only (N-(1-(pyridin-4-yl)piperidin-4-yl)acetamide) **5** used for calculations.

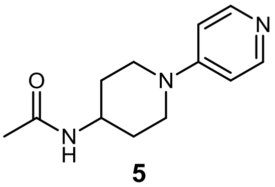

**5**

#### Free Energy Profile

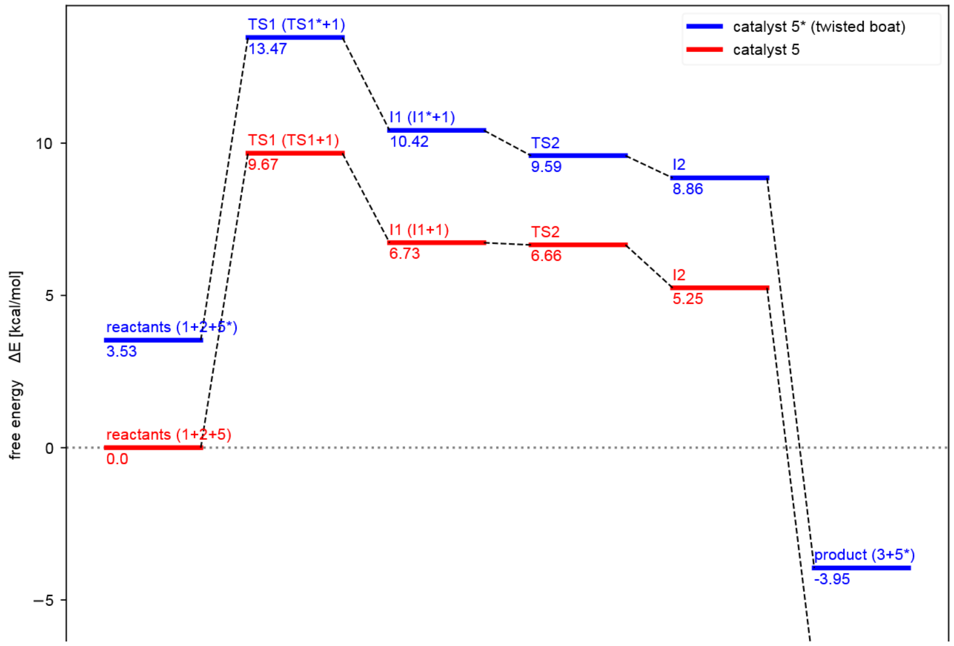

*Figure S4* Free energy profile of the Baylis-Hillman reaction between nitrobenzaldehyde **1** and cyclopentenone **2** and N-(1-(pyridin-4-yl)piperidin-4-yl)acetamide **5** as catalyst calculated at the B3LYP/ma-def2-SVP/D3BJ-CPCM // B3LYP/ma-def2-TZVP/D3BJ-CPCM level of theory.

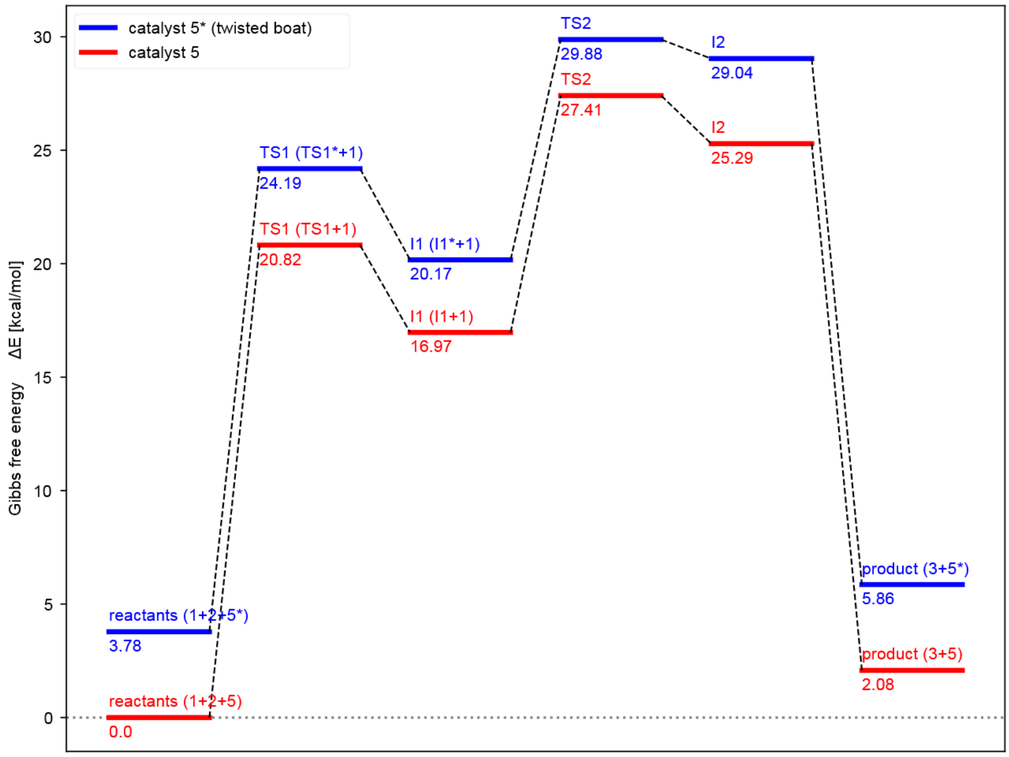

*Figure S5 Gibbs f*ree energy (entropy corrected) profile of the Baylis-Hillman reaction between nitrobenzaldehyde **1** and cyclopentenone **2** and N-(1-(pyridin-4-yl)piperidin-4-yl)acetamide **5** as catalyst calculated at the B3LYP/ma-def2-SVP/D3BJ-CPCM level of theory.

#### Exemplary input files

Final geometry optimization:

```
!B3LYP ma-def2-SVP CPCM(water) NORI Grid7 NoFinalGrid TightOpt NumFreq D3BJ moread
%scf
Convergence VERYTIGHT  # or Extreme etc.
end
%pal nprocs 28 end
%cpcm
ndiv 6
surfacetype vdw_gaussian
end
%moinp "product_final.gbw"
%base "product_rep_gaussian"
%geom
inhess Read
InHessName "product_final.hess"
end
*xyzfile 0 1 product_final.xyz
```

Transition state search:

```
!B3LYP ma-def2-SVP CPCM(water) RIJCOSX AutoAux D3BJ PAL16 NEB-TS
%neb
	NEB_End_XYZFile "final.xyz"
	Nimages 24
	SpringType DOF
	PerpSpring cosTan
	Tol_MaxF_CI 2.e-3
	Tol_RMSF_CI 1.e-3
	Tol_Scale 10.0
	Local true
	LBFGS_Mem 20
end
*xyzfile 0 1 preTS_I1.xyz
```

Final transition state optimization:

```
!B3LYP ma-def2-SVP CPCM(water) NORI Grid7 NoFinalGrid OptTS NumFreq D3BJ moread
%scf
Convergence VERYTIGHT  # or Extreme etc.
end
%pal nprocs 28 end
%cpcm
ndiv 6
surfacetype vdw_gaussian
end
%moinp "B3LYP_TS1_equi_final.gbw"
%base "B3LYP_TS1_equi_final_rep_gaussian"
%geom
inhess Read
InHessName "B3LYP_TS1_equi_final.hess"
end
*xyzfile 0 1 B3LYP_TS1_equi_final.xyz
```

#### Coordinates:

Energies given here are calculated at B3LYP D3BJ ma-def2SVP CPCM level. Geometries were the catalyst posses a twisted boat conformation are marked with a star(*).

##### Cyclopentenone

```
12
Cyclopentenone - free energy: -268.92821687 Eh, Gibbs free energy -268.96191860 Eh
 C   -0.13653903258031    -2.18142014328259     0.01659522152011
 C   -0.68139904829309    -0.94662854444867     0.03693377182522
 C    1.36581626548675    -2.16204830644799    -0.04922262461036
 C    0.38555299583508     0.06788523640079    -0.01247018757559
 C    1.72317333027628    -0.66503278308153    -0.06890839940979
 H   -0.70609053487574    -3.11512745253081     0.04399166827388
 H   -1.74341260740129    -0.69783814085483     0.08261605480559
```

| | | | |
|---|---|---|---|
| O | 0.24008140019620 | 1.28568406968991 | -0.00944613425303 |
| H | 2.34183247758879 | -0.35588855536071 | 0.78928883086457 |
| H | 2.26475807556503 | -0.36057330772655 | -0.97931072594024 |
| H | 1.72013926802931 | -2.70015239807169 | -0.94559623637993 |
| H | 1.79713741017301 | -2.69524967428539 | 0.81578876087959 |

#### *p*-Nitrobenzaldehyde

16

nitrobenzaldehyde - free energy: - 549.32676411 Eh, Gibbs free energy: -549.36987726 Eh

| | | | |
|---|---|---|---|
| C | -2.88810622256096 | 0.68096075846759 | 0.19806335132601 |
| C | -1.50513295051960 | 0.52562393641089 | 0.35265499370747 |
| C | -3.50636044753096 | 1.93286177826933 | 0.17113263641394 |
| C | -2.70638445281954 | 3.06730231744982 | 0.30427513678062 |
| C | -1.31661236281627 | 2.93967103453372 | 0.45903447444593 |
| C | -0.71998305483966 | 1.66623268488802 | 0.48300497536769 |
| H | -1.06286209043022 | -0.46989023724732 | 0.36826359738559 |
| H | 0.36144993065175 | 1.57947053251557 | 0.60418169628083 |
| H | -4.58583814321020 | 2.01306046652192 | 0.04874041001561 |
| H | -3.16322264991091 | 4.05998012151756 | 0.28700681868569 |
| C | -0.49480192977038 | 4.16877529883544 | 0.59446567114736 |
| O | 0.71462945951168 | 4.17212984351319 | 0.73040814854922 |
| H | -1.06828207688803 | 5.12378299586101 | 0.56336898395777 |
| N | -3.72297595033594 | -0.52343015715062 | 0.05686383046925 |
| O | -4.92866853366679 | -0.37621054496132 | -0.09812919993285 |
| O | -3.17189852486399 | -1.61602082942482 | 0.10107447539987 |

#### Product

28

Product - free energy: -818.27279926 Eh, Gibbs free enthalpy -818.32847663 Eh

| | | | |
|---|---|---|---|
| C | 1.45342265642328 | -1.26937638441254 | -2.30658746967904 |
| C | 1.57038982625382 | -2.79261302643448 | -2.11819747057582 |
| C | 0.46830547805579 | -3.13707403073293 | -1.15725488006513 |
| C | -0.25343859332993 | -2.06287380354461 | -0.76955718576747 |
| C | 0.29302854859816 | -0.85400381908988 | -1.41987259874472 |
| H | 1.23582419451542 | -0.97732161721981 | -3.34706658280410 |
| H | 2.35955824074204 | -0.72057804261982 | -2.00311481458913 |
| O | -0.12068347410385 | 0.29059584376625 | -1.24804008261933 |
| H | 2.54807508043558 | -3.09776642500806 | -1.70679924053614 |
| H | 1.44817924556884 | -3.35237856129619 | -3.06135490413376 |
| C | -1.45300907279416 | -1.96884892527751 | 0.14913092869290 |
| O | -1.29112335346525 | -0.89611522697097 | 1.07423004203394 |
| C | -1.68622751038365 | -3.25943679080595 | 0.90034227744397 |
| H | -2.34022195633609 | -1.77452565315792 | -0.48462117071136 |
| C | -2.51435676197403 | -4.24742222866191 | 0.34602301027031 |
| C | -1.03549034873677 | -3.50648067839576 | 2.11898963718021 |
| C | -1.20767113091033 | -4.72013401630943 | 2.78104543307319 |

C -2.69417957903010 -5.46948925437701 0.99044768377732
C -2.03575406914743 -5.68906080524229 2.20452469781316
H -0.39109154985455 -2.74043872856218 2.55213144071975
H -3.02512069580482 -4.05850151959728 -0.60116437474437
H -3.33697578798189 -6.23990932866810 0.56591985380195
H -0.70762798644178 -4.91878514192547 3.72856337762002
N -2.22113322336146 -6.96716582904006 2.89198580305314
O -2.95328327364907 -7.80615675933184 2.37613475166552
O -1.63612615341808 -7.14799486364434 3.95549335482225
H 0.28475666969783 -4.16198993602010 -0.82496133424571
H -1.04995541956756 -0.11177444741975 0.54966981724843

#### Catalyst (N-(1-(pyridin-4-yl)piperidin-4-yl)acetamide)

33
catalyst - free energy: -705.88360998 Eh, Gibbs free enthalpy: -705.94011306 Eh
C -2.45327464057337 2.20053034698472 -0.19844891586701
C -1.93062529438020 0.92678487175372 0.00306115519104
C -0.52781140003889 0.71191127101968 -0.07355191241646
C 0.23999378053708 1.87963883987237 -0.33449193692300
C -0.39298057555761 3.10367182064379 -0.51878842411619
N -1.72186204642210 3.29549792111351 -0.46039919075090
H -2.62586417705467 0.12027765279795 0.22583574126925
H -3.53878349166035 2.34015164418417 -0.13519547210093
H 1.32524687005110 1.85095726809826 -0.40855923743451
H 0.22052208577993 3.98784795207217 -0.72738678257620
N 0.04234879466922 -0.53279967455329 0.07175872692384
C -0.77326683236766 -1.68268390036795 0.46576311076644
C 1.46143824051904 -0.67181059073671 0.41221360106349
C -0.20713119964246 -3.00027040620127 -0.06282759113319
H -1.78748225439918 -1.56681911776042 0.07154960150117
H -0.85096784243641 -1.72405847570234 1.57151195779952
C 1.25576101309781 -3.18490614453848 0.33325741017173
H -0.28949463897582 -3.01634632275221 -1.16405972738228
H -0.81778067245564 -3.82930948310938 0.32848208983323
C 2.05971306867185 -1.96705783111021 -0.13099133268290
H 2.06725033060387 -1.93876176225731 -1.23422512758517
H 3.10412450432382 -2.04754702870405 0.20905394006519
H 1.57337580592915 -0.64443182302352 1.51561886921154
H 2.02820238952849 0.17193805537565 0.00959703729439
N 1.78478757899373 -4.43008715769035 -0.20395051099818
H 1.32726425718457 -3.25737973737683 1.42982112023056
C 2.66584571071893 -5.22338326717853 0.44648223340700
O 3.07194375004592 -4.97431017553640 1.59028184577167
C 3.13332628355654 -6.45422084080602 -0.30057674297085
H 2.66179261240976 -6.56662468999824 -1.28692847602427
H 2.91432720430953 -7.34435845897563 0.30925389128008
H 4.22589743506758 -6.39765583835214 -0.42733065701261

H 1.53715334996647 -4.67205491718470 -1.15896029380551
H 1.57841294836466 -4.63347533333148 -1.16496248122392

#### Catalyst* boat-conformation (N-(1-(pyridin-4-yl)piperidin-4-yl)acetamide)

33
cat_boat – free energy: -706.93063661 Eh, Gibbs free enthalpy: -705.93409623
N 12.65174655958251 2.24299400789628 -1.89403733494118
C 13.53861548717726 2.58056973893924 -0.93991348323527
C 13.22161379441300 3.24526370029641 0.23786850379637
C 11.87453040787211 3.62219048143835 0.49718127313823
C 10.93977892867024 3.25615923012347 -0.51162558738505
C 11.37669082733690 2.58965235453829 -1.65148053402159
N 11.48180167810529 4.29167621155028 1.61915279073373
C 10.06673997146807 4.62717714967506 1.81572718655722
C 12.36107917367461 4.56276573949716 2.75339097575701
C 9.86789332809663 5.50300409631769 3.05274208223361
H 9.45848508879434 3.70726672131531 1.89596843966121
H 9.70650785290690 5.17682908267083 0.92979098601962
C 10.90630919215186 6.63213741322825 3.06129121627084
C 12.33286608011036 6.04573063344131 3.14634822387470
H 13.38551043568578 4.26515922184613 2.51660156204507
H 12.04496730075382 3.93934866777217 3.60805019301305
N 10.67915767584110 7.59133056749891 4.13273607530375
H 13.01286357648500 6.62442443177068 2.50226429261679
H 12.71318239647207 6.12985438873226 4.17750939769688
H 10.79480744140256 7.18859854322567 2.11902138866922
H 8.85110754640595 5.92116762904931 3.02475069575636
H 9.95160421277693 4.91112448755743 3.97821434744723
H 14.58289759339558 2.30264342303754 -1.12358502533664
H 14.02538672481252 3.46649908866307 0.93800452240000
H 9.87913546065415 3.47980723943770 -0.41393488842705
C 9.88249227831811 8.67646521955999 4.01957749508540
H 11.07705826367939 7.38543574885177 5.04358673127307
O 9.30843027065438 8.97449265557617 2.96159004697267
C 9.73005983209714 9.52522358678513 5.26236902311388
H 10.30850219125593 9.14632650558436 6.11654855681234
H 8.66419268185891 9.56363145869368 5.53638053457566
H 10.05323056306203 10.55247892477734 5.03290888431271
H 10.64129518402840 2.31633165065262 -2.41721857178991

## TS1

45
TS1 - free energy: -974.80072459 Eh, Gibbs free enthalpy: -974.86885128 frequency: -285.44 cm**-1
C -4.32794948061295 2.35711604244078 -1.59526964633727
C -4.61769731551503 0.85145977119048 -1.49724642983134
C -4.60665321549210 0.57980723988810 -0.00038073228703

| | | | |
|---|---|---|---|
| C | -4.81816260354680 | 1.79479120992424 | 0.68068238216742 |
| C | -4.65979802366051 | 2.90099302791150 | -0.19853538897576 |
| N | -2.76808568346540 | -0.01921835710795 | 0.27868203940424 |
| H | -5.00162328465702 | -0.37109643959620 | 0.36699648496873 |
| C | -1.89132606273356 | 0.82507874985491 | 0.82555715805639 |
| C | -2.34960549920798 | -1.23479347656005 | -0.09889956742428 |
| C | -0.55980868306079 | 0.50134799078579 | 1.02351325458667 |
| C | -1.04047372943604 | -1.65478813723807 | 0.05500590050155 |
| C | -0.07611905415105 | -0.78535365431900 | 0.64630725339575 |
| H | -0.78021631298146 | -2.65567744430609 | -0.28122752520151 |
| N | 1.21736188141369 | -1.16934017933752 | 0.85488208279710 |
| C | 1.79613396246789 | -2.35077964416137 | 0.21082720312689 |
| C | 2.50813743197633 | -1.95429352573320 | -1.08780939531290 |
| H | 1.03394798095651 | -3.11378277113086 | 0.02654967458910 |
| H | 2.52073861377509 | -2.79072107233118 | 0.91538108785036 |
| C | 2.99666092045956 | 0.29043633068323 | -0.01230108622372 |
| C | 3.57797728969002 | -0.88878299672441 | -0.81149845509686 |
| H | 1.76474395185271 | -1.55992225450624 | -1.80301228248934 |
| H | 2.97303402456049 | -2.84191185434522 | -1.54591821231421 |
| H | -4.90949784020724 | 2.87652282644329 | -2.37307573682114 |
| H | -3.26029718060273 | 2.55336589928030 | -1.80042266444803 |
| H | -3.09924898704042 | -1.89469492919383 | -0.54766885584924 |
| H | 0.09046284850976 | 1.25090081885932 | 1.46849988256108 |
| H | -2.29151665851161 | 1.80521353056730 | 1.11046615549382 |
| H | -3.91429720908444 | 0.21898623873149 | -2.05806982016129 |
| H | -5.62926362138616 | 0.62276442476811 | -1.87545056785892 |
| H | -5.02410779232008 | 1.89015222939255 | 1.74887626036276 |
| O | -4.75568679112724 | 4.12524574301565 | 0.05192662831556 |
| N | 4.19737408843748 | -0.44470319215087 | -2.05120335978806 |
| H | 4.37920096585655 | -1.34684895578948 | -0.21155028862396 |
| C | 5.52683977069717 | -0.25549662596378 | -2.21392029095424 |
| C | 5.96610625009925 | 0.25112532434009 | -3.57091042172524 |
| H | 5.13004837212064 | 0.40066578266058 | -4.26834944231476 |
| H | 6.67590242312216 | -0.46960173611011 | -4.00599442843070 |
| H | 6.49994851214402 | 1.20490620493736 | -3.43682410966995 |
| C | 2.26275565359191 | -0.21598646978105 | 1.23402825056734 |
| H | 2.29300718146137 | 0.85522967428997 | -0.64783502921684 |
| H | 3.80652729669720 | 0.97725572216587 | 0.28096412653391 |
| O | 6.35106809086253 | -0.48400863879671 | -1.31774537631825 |
| H | 3.58158222272133 | -0.19369508916566 | -2.81907896045035 |
| H | 1.84813450853965 | 0.61159122990842 | 1.81746876763071 |
| H | 2.96840078678735 | -0.74590856769045 | 1.89486348121581 |

#### TS1* – boat conformation

45
TS1_equi_boat - free energy: -974.79480713 Eh, Gibbs free enthalpy: -974.86348472 Eh, frequency -277.86 cm**-1

| | | | |
|---|---|---|---|
| C | 12.92854124377125 | 0.04312669926643 | -3.23573997795350 |
| N | 12.76554137793452 | 2.46287487825825 | -2.01217831699180 |
| C | 13.66880628426576 | 2.79926060999522 | -1.08722822508358 |
| C | 13.34238311719359 | 3.48754259063172 | 0.06783121182941 |
| C | 11.99013841845522 | 3.87786428535403 | 0.30236596146217 |
| C | 11.05277431772007 | 3.49900432085982 | -0.70735567016085 |
| C | 11.48453469677592 | 2.80557767060679 | -1.82485896722781 |
| N | 11.59062658092773 | 4.56490744377576 | 1.40334913592849 |
| C | 10.16189085209740 | 4.83958680135919 | 1.61715857313420 |
| C | 12.49644878245209 | 4.97111795951993 | 2.47793424839853 |
| C | 14.19411227353600 | -0.58307490386526 | -2.62870278291299 |
| C | 15.33664046598400 | 0.32292074526357 | -3.10874172957845 |
| C | 14.76415594084592 | 1.49733608068541 | -3.66992791685109 |
| C | 13.35916451091610 | 1.45897747053682 | -3.58812145824158 |
| O | 16.54641782778864 | 0.01819906241547 | -2.99048751602106 |
| H | 15.34933369274949 | 2.33411134488843 | -4.05730459432863 |
| C | 9.92037490252630 | 5.55647167761740 | 2.94146965044826 |
| H | 9.59250037957466 | 3.89463503440770 | 1.59281109988186 |
| H | 9.78574261836726 | 5.46840543079381 | 0.79056930279716 |
| C | 10.79932222900459 | 6.80986044444811 | 3.03954962098774 |
| C | 12.29300582645437 | 6.44069480004463 | 2.85988668585691 |
| H | 13.53337125729907 | 4.82907131191934 | 2.16352299327770 |
| H | 12.33894955716590 | 4.31981494667969 | 3.35501503154525 |
| N | 10.58112172150477 | 7.53277632565097 | 4.28479994017341 |
| H | 12.74554913345117 | 7.08598607556299 | 2.09144868123519 |
| H | 12.84544499361740 | 6.62311594311684 | 3.79543161972650 |
| H | 10.49326580743988 | 7.49115077372199 | 2.23283435508014 |
| H | 8.85803001564951 | 5.83642548930291 | 2.99616901759798 |
| H | 10.12382162764973 | 4.88639410500975 | 3.79202939022067 |
| H | 14.70002669088631 | 2.49401551951863 | -1.29976687587236 |
| H | 14.13750391719861 | 3.71069472497960 | 0.77605641763570 |
| H | 9.99682779604748 | 3.74714602074285 | -0.62680481593164 |
| C | 9.67298075919749 | 8.52385883051602 | 4.43349209611985 |
| H | 11.07625579971397 | 7.21003990195735 | 5.11065403818838 |
| O | 8.97490785917486 | 8.93524681089634 | 3.49629133210371 |
| C | 9.55602578927414 | 9.12112883119117 | 5.81955963032187 |
| H | 10.25740656149850 | 8.68068918370532 | 6.54211080855452 |
| H | 8.52627000393536 | 8.97526037621827 | 6.18185555110925 |
| H | 9.73586403871168 | 10.20545784036011 | 5.75477217370849 |
| H | 12.71990153836471 | 2.06021776879587 | -4.23995728426050 |
| H | 14.17143149396417 | -0.55974182493072 | -1.52450266295190 |
| H | 14.36451213702720 | -1.62918387866486 | -2.92805797127440 |

H 12.04867587190783 0.01050543848987 -2.57682711599864
H 12.64909710277234 -0.46971469344103 -4.17256770537417
H 10.77782218720384 2.50939373183644 -2.60717698030853

## I1

45
I1 - free energy: -974.80746635 Eh, Gibbs free enthalpy: - 974.80652214 Eh
C 8.07359222737479 8.83631769616179 -4.24343620487878
N 9.14268707661353 7.81056915134211 -2.19773387377644
C 10.48016926513950 7.68431581816819 -2.33961229913331
C 11.15779635739503 6.55427939795389 -1.94779422964167
C 10.46261032500530 5.45365512328739 -1.35840519689580
C 9.04827959766845 5.62427774748760 -1.23675747756127
C 8.44512390047235 6.78477337463540 -1.65736678305022
N 11.09696437635869 4.33725343326433 -0.93211639045021
C 10.38848759473239 3.11947956618540 -0.52660940690233
C 12.50539617500113 4.05578356854840 -1.22720716493608
C 9.14580956614057 9.63731577553696 -4.99557318563576
C 9.65389130038487 10.66172826770045 -3.96493880447836
C 9.19497065734926 10.29054541888164 -2.70569260167558
C 8.39544003671310 9.04400202921441 -2.74095161052672
O 10.38893969847290 11.64535045936944 -4.31251188404089
H 9.44073286572798 10.81486516937806 -1.77801141934324
C 10.39138462220218 2.10172237394656 -1.67206763322873
H 9.36898540168983 3.34514678496276 -0.20212558716281
H 10.91762245108994 2.70519825353198 0.34684825305229
C 11.82681048480674 1.79195616207960 -2.11619401584399
C 12.60779662130550 3.08507745533853 -2.40771180033582
H 12.94673725062507 3.60181322678451 -0.32504405780243
H 13.06014777050723 4.97790249381894 -1.42118281141943
N 11.82747849244246 0.89236533693596 -3.25986778705602
H 13.66536826267282 2.84728586093607 -2.60241041105241
H 12.19951665007073 3.56584174004583 -3.31311857957846
H 12.33909922734386 1.26016773933587 -1.29939980527199
H 9.81716374390594 2.51218852276649 -2.52128104610712
H 9.89034533536289 1.17655105371045 -1.34581842767882
H 10.98674521607969 8.54622136341859 -2.77594574147341
H 12.23354349604675 6.53752878817919 -2.10187784280788
H 8.41127449711474 4.85362200574627 -0.81059178239223
C 12.72056137594297 -0.10834800635624 -3.43612788020830
H 11.17233856933561 1.08566798391435 -4.01151267311840
O 13.59648233606587 -0.37684586392316 -2.60259754847123
C 12.58962286444736 -0.90018964113019 -4.71914324489710
H 11.75447968534044 -0.56581345738802 -5.35037792296223
H 12.45118422533419 -1.96321519202004 -4.46783335629552
H 13.52918564461566 -0.81160201193555 -5.28675697459749
H 7.48391933251087 9.04753423369847 -2.12412425561792

| | | | |
|---|---|---|---|
| H | 9.99240915047959 | 9.00523707369870 | -5.32260322429833 |
| H | 8.75709029813376 | 10.13904216237626 | -5.89687283483052 |
| H | 8.03430205429295 | 7.76947286758061 | -4.51393379067519 |
| H | 7.07533799067666 | 9.25984147454400 | -4.43616820089334 |
| H | 7.36632592900754 | 6.92870721828754 | -1.57351048404841 |

#### I1* boat

45

I1 - free energy: -974.80169440 Eh, Gibbs free enthalpy: -974.86988627 Eh

| | | | |
|---|---|---|---|
| C | 13.04368032839888 | 0.17657849872387 | -3.00364475887218 |
| N | 12.66534261408659 | 2.45194889847812 | -1.97540137744615 |
| C | 13.57254115912395 | 2.72153115411837 | -1.00914970691693 |
| C | 13.22379604673923 | 3.30298023388782 | 0.18582106044988 |
| C | 11.86500937641680 | 3.65790079854645 | 0.45159481239324 |
| C | 10.93674755607574 | 3.35456587522764 | -0.59437073672211 |
| C | 11.36717363280787 | 2.76312611512512 | -1.75804337874397 |
| N | 11.45892308475862 | 4.24369147516596 | 1.59624116722384 |
| C | 10.04531017020272 | 4.61514057933825 | 1.78289312312497 |
| C | 12.32414114735447 | 4.46926077798570 | 2.75623310075534 |
| C | 14.50960892242494 | -0.21177400497264 | -2.76363186687147 |
| C | 15.31902073221371 | 0.92227342536007 | -3.41809064578417 |
| C | 14.45923170578947 | 1.97876356602595 | -3.69725800011086 |
| C | 13.06986169252033 | 1.70516286367573 | -3.26076602527511 |
| O | 16.57714413150281 | 0.81765382195735 | -3.60844005553218 |
| H | 14.77278506618313 | 2.93696707822725 | -4.12072480962028 |
| C | 9.85609660331064 | 5.47991143006416 | 3.02855678675083 |
| H | 9.41975111289482 | 3.70708708630938 | 1.83974710689446 |
| H | 9.71869282839720 | 5.18853649402280 | 0.90079145141569 |
| C | 10.92814635304599 | 6.57738539575603 | 3.06273773888134 |
| C | 12.33349147105836 | 5.94525671951306 | 3.17284282797479 |
| H | 13.34142971249750 | 4.13295495091345 | 2.54446235032963 |
| H | 11.95299911909041 | 3.84211499533611 | 3.58309356151884 |
| N | 10.70718370017576 | 7.54161366712577 | 4.12959884849185 |
| H | 13.04919764494510 | 6.51253121532862 | 2.55846753395984 |
| H | 12.68878272097256 | 5.99508742600234 | 4.21488255707393 |
| H | 10.85315204954791 | 7.13946667045459 | 2.12038338600475 |
| H | 8.85268463043350 | 5.92806065804698 | 2.98749058911867 |
| H | 9.90637273263899 | 4.87582416736929 | 3.94840889832836 |
| H | 14.60329896744781 | 2.46092631955202 | -1.25231718306687 |
| H | 14.01865771484635 | 3.49055866864554 | 0.90371376235981 |
| H | 9.87618928126962 | 3.57263980050602 | -0.49740753596859 |
| C | 9.97524593806597 | 8.67032756865168 | 3.98202933444857 |
| H | 11.02916426241257 | 7.29792853557258 | 5.06162552330447 |
| O | 9.47959665806783 | 9.00501697030839 | 2.89740013638632 |
| C | 9.79418091123988 | 9.51741781664203 | 5.22290935801500 |
| H | 10.34281470525176 | 9.13130326905365 | 6.09331035598386 |
| H | 8.72068662147394 | 9.56298363861609 | 5.46548898541481 |

| | | | |
|---|---|---|---|
| H | 10.13120005926066 | 10.54269463697313 | 5.00623535608821 |
| H | 12.28206066879916 | 2.01578373269254 | -3.96369140246699 |
| H | 14.76213952018652 | -0.26011435900913 | -1.68797913450009 |
| H | 14.77354606271746 | -1.19065552330999 | -3.19641288922401 |
| H | 12.36448055870155 | -0.09299195190534 | -2.17969541433596 |
| H | 12.66394307413502 | -0.31190761450645 | -3.91480872801887 |
| H | 10.67201695051172 | 2.51563645840229 | -2.56246606321476 |

## TS2

61

TS2 - free energy: -1524.13973596 Eh, Gibbs free energy -1524.22823218 Eh, frequency -328.87 cm**-1

| | | | |
|---|---|---|---|
| C | -2.93243815064824 | 2.40437022144019 | -1.98388914164598 |
| C | -2.82569188574410 | 0.87653018458752 | -1.86454666373874 |
| C | -2.67898266886101 | 0.59122783650583 | -0.34918895202920 |
| C | -3.38318426400563 | 1.74775368170496 | 0.31381923094101 |
| C | -3.29637988149805 | 2.89166128610699 | -0.57990567119223 |
| N | -1.23649875792702 | 0.45183125327403 | 0.05169062082961 |
| H | -3.10645909148373 | -0.38370964810287 | -0.08981993603179 |
| C | -0.43772140872029 | 1.52432159607632 | 0.27892353000817 |
| C | -0.68575985320909 | -0.78854204188851 | 0.10032122970483 |
| C | 0.89949576360020 | 1.39695826153916 | 0.55909762049265 |
| C | 0.64142790540902 | -0.99521320826181 | 0.37613314060182 |
| C | 1.51580443718029 | 0.10839836614575 | 0.62891692367410 |
| H | 0.99204144105567 | -2.02389129949644 | 0.39374616937536 |
| N | 2.82234114227014 | -0.05545427623490 | 0.92331266442778 |
| C | 3.77502494936190 | 1.05760560821390 | 0.99320865319185 |
| C | 3.51930897549003 | -1.33966998738663 | 0.79221633349813 |
| C | 4.30581868950451 | -1.37000278360325 | -0.52175354722026 |
| H | 2.81946761255632 | -2.17705394560369 | 0.85735814814851 |
| H | 4.20646161475881 | -1.42989743435941 | 1.64886042118942 |
| C | 4.57885828339929 | 1.14163229040774 | -0.30819892417962 |
| H | 4.44881995148034 | 0.85954766717797 | 1.84209465141641 |
| H | 3.26470792690154 | 2.00106487394435 | 1.20546480863212 |
| C | 5.28857509798319 | -0.18853304554932 | -0.59432616859799 |
| H | 3.60207272277404 | -1.31735805378932 | -1.37006056014378 |
| H | 4.85694196691555 | -2.32007173119830 | -0.60200526669239 |
| H | 5.31878956966649 | 1.95405505569294 | -0.23299303130673 |
| H | 3.89518089013687 | 1.38723236428241 | -1.13971182334431 |
| N | 5.97423743491209 | -0.13740704108117 | -1.87634485235502 |
| H | 6.06576063111944 | -0.34276300145009 | 0.17002457068216 |
| C | 7.20161111148073 | -0.65967141562642 | -2.10298912091453 |
| C | 7.72262695335501 | -0.54533795603241 | -3.51932040731256 |
| H | 8.68534883577358 | -0.01141441150773 | -3.50129803614591 |
| H | 7.03099751072475 | -0.02180959918555 | -4.19409030971905 |
| H | 7.91055735621862 | -1.55780608239710 | -3.90984567047901 |
| H | 5.46312288942602 | 0.24435794397748 | -2.66673298126250 |

| | | | |
|---|---|---|---|
| O | 7.87427456326606 | -1.19756818195155 | -1.21299856357381 |
| H | -3.68155334452453 | 2.74023361555723 | -2.71875850486176 |
| H | -1.97456735474739 | 2.87393280424130 | -2.26737303076609 |
| O | -3.50069656621355 | 4.07750830468967 | -0.29692638688723 |
| H | -1.36176315015136 | -1.62173818624403 | -0.09615279841909 |
| H | 1.46223899102946 | 2.31207895434451 | 0.72294445331640 |
| H | -0.90904153536381 | 2.50470486707839 | 0.23697832094603 |
| H | -1.99724813770230 | 0.44089315095393 | -2.44126823017127 |
| H | -3.75400127050848 | 0.39380959280183 | -2.20546846218886 |
| C | -5.33816339455666 | 1.44148540335594 | 0.37394858242169 |
| H | -3.22674672546428 | 1.91574465844648 | 1.38346537531647 |
| O | -5.92410904369050 | 2.44356270168282 | 0.90927769870775 |
| C | -5.30818246141179 | 0.13101025216351 | 1.12813883375507 |
| H | -5.49648494966599 | 1.25686114868833 | -0.71347114165543 |
| C | -5.40285429229747 | -1.09010602572350 | 0.43885996659935 |
| C | -5.20016222290502 | 0.11701517807355 | 2.53072060912105 |
| C | -5.15897773064152 | -1.08251514438369 | 3.23112855881751 |
| C | -5.37026976588785 | -2.30409795455032 | 1.12214395324048 |
| C | -5.24359673805448 | -2.28644856141535 | 2.51536032589507 |
| H | -5.14032674827917 | 1.06517065949735 | 3.06860997323896 |
| H | -5.50410170910680 | -1.08837896790595 | -0.64947724432640 |
| H | -5.44228389478184 | -3.25203000415261 | 0.58964728649889 |
| H | -5.06277299379205 | -1.09804923037704 | 4.31649521084687 |
| N | -5.19967236296392 | -3.54936694518697 | 3.24036497187039 |
| O | -5.28544262976328 | -4.59680895135977 | 2.60254978201417 |
| O | -5.07578023317875 | -3.51984966664675 | 4.46309380774140 |

#### TS2* – boat conformation

61

TS2 – free energy: -1524.13578945 , Gibbs free energy -1524.22429719 Eh, frequency -343.11 cm**-1

| | | | |
|---|---|---|---|
| C | 0.52429508241139 | -2.75583686159224 | -1.94441241536668 |
| N | 0.00620613618154 | -0.49107349160245 | -0.94630816666737 |
| C | 1.04661987852020 | -0.23311346636467 | -0.11393649035448 |
| C | 0.88334127280796 | 0.41659199126213 | 1.08277961713205 |
| C | -0.41195055277763 | 0.86187601515027 | 1.50249005907385 |
| C | -1.48262033694087 | 0.56114273484282 | 0.60345477460265 |
| C | -1.23934682734179 | -0.09878984052421 | -0.57472930818246 |
| N | -0.58015346426928 | 1.51219359599774 | 2.66789768583625 |
| C | -1.87020334038418 | 2.04082989016197 | 3.12410939859842 |
| C | 0.55924137532177 | 1.68669473907986 | 3.58819755386243 |
| C | 2.02744453322096 | -2.93403870254243 | -2.21710543722516 |
| C | 2.49015811664837 | -1.63453669966238 | -2.87300031588641 |
| C | 1.35867705377207 | -0.74320130812267 | -3.04103218201117 |
| C | 0.20407070260374 | -1.25798905659534 | -2.21873634455897 |
| O | 3.66001141627357 | -1.42424850469405 | -3.21942623623023 |
| C | -1.73689093166036 | 3.48600780902188 | 3.61026727343705 |

| | | | |
|---|---|---|---|
| H | -2.26096591276977 | 1.39335769806378 | 3.92638617474212 |
| H | -2.59322526186208 | 2.00789206112831 | 2.30540533326288 |
| C | -0.57802910262190 | 3.67845892332075 | 4.62163457218889 |
| C | 0.12769240214752 | 2.34316692942587 | 4.89364271379050 |
| H | 1.32673245994204 | 2.31047952139429 | 3.09743256744933 |
| H | 1.01324593734904 | 0.70419294668421 | 3.79413462323776 |
| N | -1.04166614189948 | 4.30312903919701 | 5.85240257683326 |
| H | 1.02355053792040 | 2.51075673530969 | 5.50928905108429 |
| H | -0.53636400721349 | 1.67348385808685 | 5.46302167050131 |
| H | 0.16512179794109 | 4.36513813368988 | 4.19326786352224 |
| H | -1.58838782511792 | 4.14751165662591 | 2.74327546196105 |
| H | -2.69520569938200 | 3.77349685127797 | 4.06990530160700 |
| H | 2.03132320052350 | -0.55971456528218 | -0.44568308785032 |
| H | 1.76995757974644 | 0.58904063819026 | 1.68754318248300 |
| H | -2.51367189752558 | 0.82846979054564 | 0.82137965783391 |
| C | -0.32128823262099 | 5.20513275394952 | 6.55819317349050 |
| H | -1.91706541493098 | 3.97539312336274 | 6.24989600413927 |
| O | 0.78598661570681 | 5.61534417951142 | 6.18425195450856 |
| C | -0.94786110835200 | 5.69902509819630 | 7.84421672882684 |
| H | -1.95890735629327 | 5.30276299501185 | 8.01266251706524 |
| H | -0.98681884791945 | 6.79888591058440 | 7.82149503884687 |
| H | -0.30183021014399 | 5.40505623411330 | 8.68659122868250 |
| H | -0.75894313768648 | -1.14900558488249 | -2.73565938770411 |
| H | 2.61557476901834 | -3.07586474233534 | -1.29455918200916 |
| H | 2.25174540962024 | -3.79096815918118 | -2.87195323941793 |
| H | 0.23838955588003 | -3.03108468594573 | -0.91923714377125 |
| H | -0.07672872635750 | -3.37476763419992 | -2.62258535450519 |
| H | -2.04493437409617 | -0.34053811059919 | -1.26900794838555 |
| C | 0.89588477247890 | -0.76829520979125 | -4.96405006471327 |
| C | -0.80743099717398 | -2.63853737577471 | -5.12336570788890 |
| C | -1.15590253722675 | -3.96632395359135 | -5.35694738621817 |
| C | -0.14764940016272 | -4.87402372722652 | -5.70160732079965 |
| C | 1.18977402900711 | -4.47328143728276 | -5.83023371893973 |
| C | 0.52912967628687 | -2.21173356231569 | -5.22168293382162 |
| C | 1.51535782270524 | -3.14182615673905 | -5.59789223838223 |
| H | 2.54937058375557 | -2.80901245081815 | -5.69746850975232 |
| H | 1.95470323833946 | -5.19880822675283 | -6.10552932422540 |
| H | -2.19060210862547 | -4.29869143130442 | -5.27855696121952 |
| H | -1.58782139311383 | -1.91867730397054 | -4.86537914720429 |
| H | -0.02114458035635 | -0.16708855514426 | -4.76087770432668 |
| N | -0.49837665663382 | -6.26752139017205 | -5.94165136306324 |
| O | 0.38881417631349 | -7.04531315916263 | -6.28718720046847 |
| O | -1.66778770415007 | -6.61400994106433 | -5.78697297327274 |
| H | 1.55061672057105 | 0.33333316075894 | -3.04245742635171 |
| O | 1.83673723459553 | -0.24092971870849 | -5.64599153782562 |

### I2

61

I2 – free energy: -1524.14229454 Eh, Gibbs free energy -1524.23161151 Eh

```
C 0.06651678377400 -0.05064711671327 -1.46354751936193
C 0.36520724703836 -1.43266264623599 -2.06431300411721
C 0.59758031432521 -2.39876477746239 -0.87516370687708
C -0.07814494662352 -1.74831408656050 0.34473428554204
C -0.15428014069315 -0.27902744138529 0.02634670534664
N 2.02540098512484 -2.70863365255389 -0.61217090639904
H 0.15279973361688 -3.37259530099833 -1.10189185742417
C 2.96532215519420 -1.73254311617717 -0.51033812158813
C 2.39830636898294 -3.99581132668368 -0.38846761774257
C 4.27263446063487 -2.00402821172952 -0.20125880876765
C 3.68606946085516 -4.34248061725890 -0.07072086737286
C 4.70516024991824 -3.34606853552795 0.04843282978061
H 3.89417142170762 -5.39899872768838 0.07493100389343
N 5.97636854693428 -3.64708631935453 0.37908798904334
C 7.08563299865474 -2.69148513267888 0.28901178420654
C 6.46939812738247 -5.02145401351011 0.52063229272541
C 7.27735990544793 -5.41844890715332 -0.71808946408534
H 5.64660084831447 -5.71842115606128 0.70042269969696
H 7.11195561648173 -5.04720604128742 1.41574753146218
C 7.92377080536871 -2.98185497356368 -0.96025603698472
H 7.70077518147265 -2.81303125938407 1.19522180849949
H 6.72083339421902 -1.66124832904122 0.28540952815690
C 8.43196493675748 -4.43019254742162 -0.95059017572395
H 6.61359171368016 -5.42810503624477 -1.59974837622020
H 7.67657075722512 -6.43666658930070 -0.58823601827590
H 8.77562395342505 -2.28492403772508 -1.00169789141953
H 7.30511568447755 -2.80934083823360 -1.85844580511526
N 9.15717035102573 -4.73282912382603 -2.17400844350883
H 9.15314049110454 -4.54282267405081 -0.12699112961694
C 10.38173648057982 -5.30696136839761 -2.21638356397942
C 10.95584849786860 -5.54667972238708 -3.59626419877141
H 11.89893571710832 -4.98542871924164 -3.68861499780207
H 10.27684146307781 -5.24489624840166 -4.40612100250062
H 11.19305523427703 -6.61663565237926 -3.70141528258730
H 8.68869048249970 -4.54021153136253 -3.05453712903744
O 11.01314442118376 -5.61657683597305 -1.19691067171484
H -0.82299980902403 0.43362254761883 -1.89837727328270
H 0.89828347902135 0.66466854190773 -1.57990935756748
O -0.34577420989038 0.61336810512071 0.83673991608660
H 1.61347813777629 -4.74706065857644 -0.47885239218421
H 4.95481818680847 -1.15979821232858 -0.14710184472747
H 2.63943407715831 -0.70943718569613 -0.69115569017682
H 1.21841680860779 -1.43474922406560 -2.75767890360529
H -0.50436475492635 -1.80437144572365 -2.62460856887971
C -1.55158136446788 -2.37318974817091 0.58652589742285
```

H 0.47765975074778 -1.90603440888175 1.27987640944606
O -2.23537893477558 -1.80022597769741 1.58605347215217
C -1.33925177799267 -3.87630944420891 0.82798605831704
H -2.05114637439306 -2.29333465699152 -0.41545382567871
C -1.90163370143911 -4.83448984441908 -0.03117826814713
C -0.60485204723439 -4.31925447806175 1.94412884772002
C -0.41135148130821 -5.67374517034777 2.18725322397171
C -1.72689696353537 -6.19919686554091 0.19467705987254
C -0.97612654796807 -6.60556192508603 1.30307194088369
H -0.17086048521767 -3.58815173461915 2.62944613655991
H -2.47654089492113 -4.50425903730510 -0.90029041203381
H -2.15654588784006 -6.94103726222522 -0.47815445159866
H 0.17175494228599 -6.01520321329816 3.04209698937232
N -0.77458290727135 -8.02683630164104 1.54310977805022
O -1.27872023788204 -8.83339244133176 0.76336760061617
O -0.10638670474258 -8.36926734447336 2.51728179605176

#### I2* – boat conformation

61
I2 – free energy: -1524.13823874 Eh, Gibbs free energy -1524.22563646 Eh
C 12.79600714175715 0.15360172787976 -3.23344296789760
N 11.80478131779220 2.08564889083523 -1.91862797786662
C 12.83334090542977 2.18894545972781 -1.03423022256209
C 12.69820303014879 2.81110477026997 0.17892197967598
C 11.45186281043845 3.39948267819040 0.56353895618800
C 10.39470504275486 3.27221105216211 -0.39297911870951
C 10.60528478283561 2.62922438437191 -1.58650533763024
N 11.26448159374334 4.03155281197039 1.73463454051421
C 9.98079208986370 4.69152331587645 2.03768340751501
C 12.26073507310435 4.09095284512044 2.80746705591220
C 14.22631066909900 0.50754070584821 -3.67058085486417
C 14.15627584093331 1.92602055851409 -4.22339555337929
C 12.73548481345360 2.41441784023394 -4.21836626527936
C 11.98478479438580 1.47311613719510 -3.26334742834887
O 15.12864509256360 2.56292774866441 -4.60153681593702
C 10.10694729332576 5.62050868132665 3.24567514277425
H 9.19648051241576 3.93308096587094 2.20490164829116
H 9.68666881521995 5.28968978130662 1.16146662273338
C 11.38749671370085 6.45819152088849 3.12069649004585
C 12.63009601750088 5.54013084863339 3.14517867669682
H 13.15237746811266 3.52179713651910 2.53531758843238
H 11.82893594663478 3.59197640523547 3.68945145718655
N 11.48428666125161 7.48972877065961 4.14107049175241
H 13.38917584653800 5.91639274043796 2.44261892036975
H 13.08444743179697 5.54375815375668 4.14929235365909
H 11.34146299190719 6.98335732441005 2.15522642451576
H 9.22407477756900 6.27572579380852 3.26849658148310

| | | | |
|---|---|---|---|
| H | 10.11621962397037 | 5.05372855314840 | 4.19022448202072 |
| H | 13.78576022135209 | 1.75228034034084 | -1.32823021294692 |
| H | 13.57493351108336 | 2.84712091249697 | 0.82055516427250 |
| H | 9.39753837437127 | 3.65933232420154 | -0.19880110591730 |
| C | 11.00351520073629 | 8.74546438476467 | 3.98864714560995 |
| H | 11.85031011389866 | 7.22429144658628 | 5.05035294270138 |
| O | 10.47386493563256 | 9.13247466090490 | 2.93827268734154 |
| C | 11.15668480476156 | 9.66767069989079 | 5.17850097694174 |
| H | 11.66187469685280 | 9.19294745208651 | 6.03123765601419 |
| H | 10.15799729551106 | 10.00820801071510 | 5.49370569915747 |
| H | 11.72868166291131 | 10.55560632070170 | 4.86734221945311 |
| H | 10.96251734779437 | 1.28821494938452 | -3.61379667083868 |
| H | 14.95196880161520 | 0.49870957464166 | -2.84009045499657 |
| H | 14.62799941272604 | -0.17015418291143 | -4.44024103248074 |
| H | 12.74634625644527 | -0.32696688217843 | -2.24618265688384 |
| H | 12.33544855546029 | -0.54064430239108 | -3.94814647937060 |
| H | 9.80803686904480 | 2.52017482637378 | -2.32249472818848 |
| C | 12.06507561171144 | 2.43989204452343 | -5.71632273266250 |
| C | 10.85994554666405 | 0.23968045979169 | -6.24033830231647 |
| C | 10.83007656810422 | -1.05838539399521 | -6.74652573404697 |
| C | 11.99683309156198 | -1.58486497194503 | -7.31126969594744 |
| C | 13.17643032299915 | -0.83087973599371 | -7.39238743387373 |
| C | 12.03211490100563 | 1.01409164681939 | -6.28525412722039 |
| C | 13.17811752863859 | 0.46522464074285 | -6.88947315259954 |
| H | 14.08416524527074 | 1.06912907382100 | -6.96025878173524 |
| H | 14.07073628377525 | -1.26202727310509 | -7.84164927183091 |
| H | 9.92148153476321 | -1.65860794908589 | -6.70502528100145 |
| H | 9.95368156690748 | 0.66048679053534 | -5.79748858730303 |
| H | 11.00442399691934 | 2.67609312015566 | -5.43693282434604 |
| N | 11.98482631930204 | -2.94627600691013 | -7.82886340605117 |
| O | 13.01969004700844 | -3.39905719560870 | -8.31436693025212 |
| O | 10.94176730224859 | -3.59338801716775 | -7.75823104607393 |
| H | 12.67633564283632 | 3.46875865949123 | -3.91473816411424 |
| O | 12.63252533183227 | 3.33005196945861 | -6.52897595578456 |

#### Protein Expression and Purification

Buffer SEC:
Potassium phosphate 50mM pH 8.0, NaCl 150 mM

Buffer IB1
TRIS* HCl (20 mM, pH 8.0), Triton-X (1 % v/v), EDTA (1 mM)
Buffer IB2:
TRIS* HCl (20 mM, pH 8.0), EDTA (1 mM), NaCl (1 M)
Buffer IB3:
TRIS* HCl (20 mM, pH 8.0), EDTA (1 mM)

Streptavidin variants were expressed using *E. coli* T7 Express lysY/Iq (New England Biolabs) with tight T7 promotor control. In baffled flasks (3 L) LB-Miller (1 L) containing ampicillin was inoculated with 10 ml preculture at 37 °C. After reaching an $OD_{600}$ of 0.8 protein expression was induced with IPTG (1 mM) and continued at 37 °C for four hours. The cells were harvested (4 400 x g, 12 min), washed with Buffer SEC and centrifuged again. Pellets were stored at -20 °C.

The protein was mainly in inclusion bodies (IB). The IB were prepared using following 4-step protocol:

1) The cells were dissolved in buffer IB1 (~10 ml/1g pellet) and lysed via sonication (3 min). The insoluble fraction was separated by centrifugation (40 000 x g, 20 min). The soluble fraction was discarded.
2) Step 1 was repeated with the pellet.
3) The insoluble fraction was resuspended in buffer IB2 to remove DNA/RNA fragments and the insoluble fraction separated again by centrifugation (40 000 x g, 20 min).
4) The insoluble fraction was resuspended in buffer IB3 to remove salt and the insoluble fraction separated again by centrifugation (40 000 x g, 20 min).

The IB was dissolved in Guanidium*HCl (6 M, pH 1.5, ~10 ml/1g pellet), centrifuged (40 000 x g, 20 min) and dialyzed against Guanidium*HCl (6 M, pH 1.5, 500-1000 ml) to lower the concentration of biotin.

The protein was then refolded in buffer SEC (2 x 1 L, for at least 10 hours each), centrifuged (40 000 x g, 20 min).

According to SDS-PAGE the protein is almost pure but was further purified using a preparative Superdex S75 column.

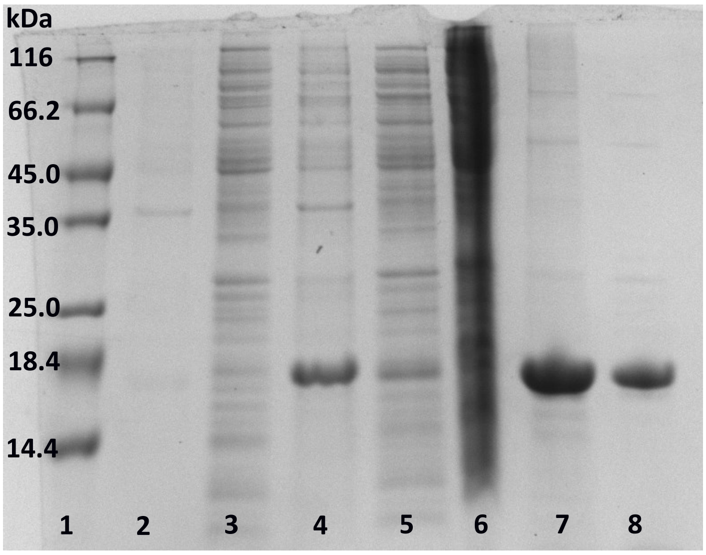

**Figure S6.** *Exemplary SDS-PAGE-Gel of streptavidin variant S112I Q114A R121A L124Y. Lane 1 : Molecular weight marker; Lane 2: Pellet before induction; Lane 3: Lysate before induction; Lane 4: Pellet after induction; Lane 5: Lysate after induction; Lane 6: Lysate of first inclusion body preparation step; Lane 7: Refolded inclusion body; Lane 8: After size-exclusion chromatography.*

#### Determination of free Biotin binding sites

The determination of biotin-free binding sites of streptavidin was done as described using Biotin-4-fluorescein (B4F, CAS 1032732-74-3 from aatbio). [20]

B4F was dissolved DMSO (11.75 mM) and 400x diluted with buffer SEC (29.4 µM). All measurements were done in duplicates on a Tecan Spark plate reader in black 96-well plates (Excitation wavelength 485 nm, Emission wavelength 520 nm, Slit 10 nm).

To a B4F solution (10 µl, 29.4 µM), streptavidin (12.9 µM) was added in a final concentration of 0 µM, 0.65 µM, 1.29 µM, 1.95 µM, 2.58 µM, 3.24 µM, 3.88 µM and 4.53 µM and buffer SEC was used to fill up to a final volume of 100 µl. All measurements were carried out as duplicates.

An exemplary graphical representation of the results is shown below. All constructs have more than 3 free biotin binding sites (between 3.2 and 3.4).

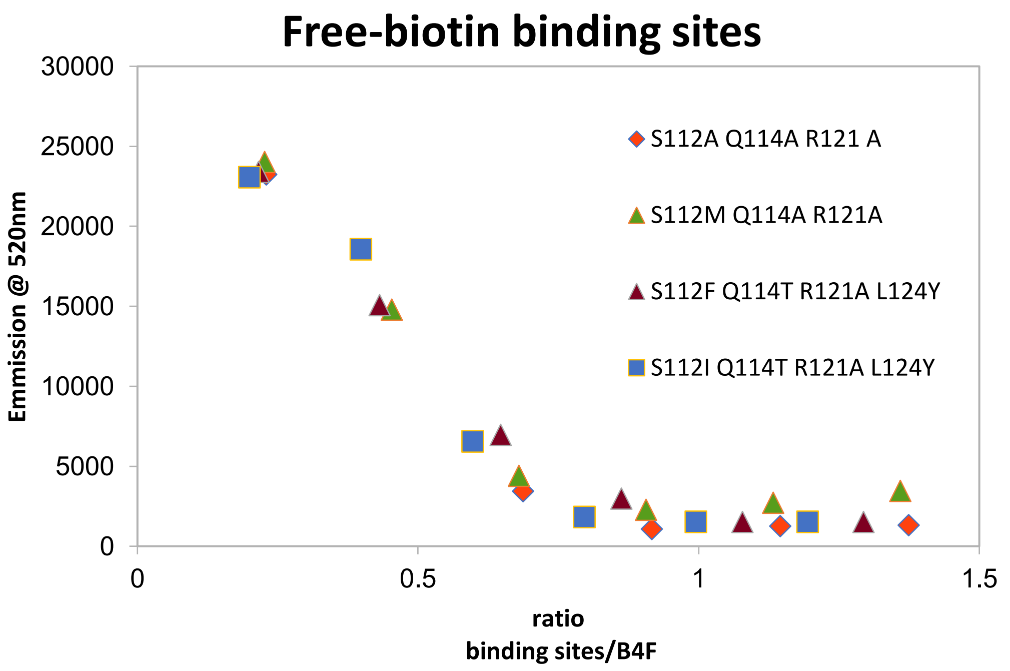

**Figure S7.** Free biotin binding sites as determined by B4F-assay in duplicates.

#### Isatin results

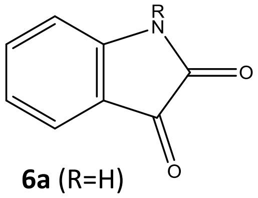

**6a** (R=H)
**6b** ($R=CH_3$)

**Table S4.** Activities and selectivities of catalysts and selected streptavidin constructs for the model substrates **5** + **2**.

| Entry | Substrate | Catalyst | Mol% Cat. | Reaction time [h] | Conv. [%][a] | HPLC area 254 nm | e.e. [%][a] |
|---|---|---|---|---|---|---|---|
| 1 | **6a+2** | **S112M** Q114A R121A L124Y | 2 | 48 | 37 | 449 | n.d. |
| 2 | **6a+2** | **S112M** Q114A R121A L124Y + **4** | 2 | 48 | 87 | 1083 | n.d. |
| 3 | **6b+2** | **S112M** Q114A R121A L124Y | 2 | 48 | | 885 | n.d. |
| 4 | **6b+2** | **S112M** Q114A R121A L124Y + **4** | 2 | 48 | | 1232 | <5 |

Reaction conditions: HEPES buffer (10 mM, pH 7.0), streptavidin (2.6 mol%), DMSO (20 vol%), isatin (50 mM), cyclopentenone (100 mM), 30 °C, orbital shaking 1000 rpm. [a] Determined by HPLC [b] 1.3 mol%. n.d. not determined

#### Reaction setup

The reactions were carried out in plastic tubes (1.5 ml) in a 50 µl-scale. To streptavidin (always assuming at least 3 free biotin-binding sites) the artificial cofactor (10 mM in DMSO) was added and incubated for 1-2 hours. Nitrobenzaldehyde/Isatin was added from a DMSO stock (500 mM). Finally cyclopentenone was supplemented. The reaction was carried out using an orbital shaker at 800-1000 rpm/30 °C). After the corresponding time the reaction was stopped by adding $NH_4CH_3COOH$ buffer (20 mM, pH 4.5, 300µl) and acetonitrile. The sample was further diluted with acetonitrile (15:85, sample: acetonitrile) and cleared by centrifugation.

#### HPLC

An Agilent Infinity 1260 HPLC system equipped with a G1316A detector was used. The compounds were separated using a Phenomenex Luna C18(2) 5µm, 250 x 4.6 mm column at 30°C. Using acetonitrile and water (+.1% TFA) as eluent following gradient was used: 1 ml/min, 10% acetonitrile for 1 min, then 10% to 70% acetonitrile over 10 minutes and final 10% acetonitrile for further 3 minutes.

Detection of nitrobenzaldehyde and the corresponding Baylis-Hillman product 2-(hydroxy(4-nitrophenyl)methyl)cyclopent-2-enone at 280 nm, while isatin and 3-hydroxy-3-(5-oxocyclopent-1-en-1-yl)indolin-2-one were detected at 254 nm.

For chiral separation a Shimadzu LC20-AD with an SPD-M20A detector was employed. On a Chiralcel® OJ column (10 µm, 250 x 4.6 mm) a isocratic elution (heptane:*iso*-propanol, 90:10) at 30 °C lead to the separation of product **3** (retention times: 36.8 min (*S*) and 43.4 min (*R*)) as reported.[21]

Products of the reaction with Isatin could not be separated using this column, but the products of the reaction N-methyl isatin were well separated (enantiomer1 11.2 min, enantiomer2 14.9 min).
